# Modeling the three-way feedback between cellular contractility, actin polymerization, and adhesion turnover resolves the contradictory effects of RhoA and Rac1 on endothelial junction dynamics

**DOI:** 10.1101/2021.03.15.435512

**Authors:** Eoin McEvoy, Tal Sneh, Emad Moeendarbary, Gloria E. Marino, Xingyu Chen, Jorge Escribano, Fabian Spill, Jose Manuel Garcia-Aznar, Ashani T. Weeraratna, Tatyana M. Svitkina, Roger D. Kamm, Vivek B. Shenoy

## Abstract

The formation and recovery of gaps in the vascular endothelium governs a wide range of physiological and pathological phenomena, from angiogenesis to atherosclerosis and tumor cell extravasation. However, the interplay between the mechanical and signaling processes that drive dynamic behavior in vascular endothelial cells is not well understood. In this study, we propose a chemo-mechanical model to investigate the maintenance of endothelial junctions as dependent on the crosstalk between actomyosin contractility, VE-cadherin bond turnover, and actin polymerization, which mediate the forces exerted on the cell-cell interface. Our theoretical model reveals that active cell tension can stabilize cadherin bonds within an adhesion, but excessive RhoA signaling can drive bond dissociation and junction failure. While Rac1-mediated actin polymerization aids gap closure, high levels of Rac1 may also facilitate junction weakening. Combining the modeling framework with novel experiments, we identify how dynamic rupture and heal cycles emerge and, further, describe why gaps tend to localize at multi-cell contacts. Beyond, our analysis also indicates that a critical balance between RhoA and Rac1 expression is required to maintain junction stability and limit endothelial dysfunction. The model predicts how pharmacological modulation of actin polymerization and cell contractility impacts junction stability, with predictions subsequently validated experimentally. Our proposed framework can help guide the development of therapeutics that target the Rho family of GTPases and downstream active mechanical processes.

## Introduction

The maintenance and turnover of endothelial junctions govern a wide range of vascular activity, from capillary barrier function to vessel branching during angiogenesis (1). Vascular homeostasis is closely tied to the regulation of this complex system, with junction breakdown mediating leukocyte migration, bruising, and edema (2, 3), in addition to pathological events such as hemorrhagic stroke (4), onset of atherosclerosis (5), and tumor cell invasion (6). As such, endothelial junctions undergo frequent dynamic rearrangements to modify their structure and porosity. Cell-cell adhesion is maintained predominantly by adherens junctions (AJs), comprised of cadherin bonds which interface with the actomyosin network and whose stability is tension-regulated (7, 8). Cadherin is therefore essential to normal endothelium activity; blocking VE-cadherin’s functions using the BV13 antibody completely disassembled blood vessels in mice (9), and cadherin knockout is also implicated in loss of barrier function (10, 11). Further, cadherin endocytosis is a mechanism by which vascular endothelial growth factor (VEGF) can form gaps in existing blood vessels to permit the growth of new capillaries (12). Unbound cadherin, as a transmembrane protein, is free to diffuse along the membrane, but typically it preferentially localizes at cell-cell contacts (13). Adhesions at these contacts stabilize and reinforce under tension (14), establishing a feedback loop with the actomyosin cables which the junctions sustain. These key cadherin interactions are also calcium dependent and strengthen under the influx of Ca^2+^ from mechanosensitive ion channels (15).

Forces acting on intercellular junctions trigger a range of biochemical processes that activate the Src family of kinases (SFK), which downstream interact with Rho-GTPases to drive changes in myosin motor activity and cell contractility (16). Upregulation in RhoA activity can initiate high engagement of myosin with actin filaments to yield dense stress fibers, reported to promote the clustering of cadherin proteins and impose extremely high bond tension (17, 18). Induction of this pathway via thrombin treatment causes poor endothelial barrier integrity and junction permeability (19–21); thus thrombin in relatively high doses is often considered a destabilizing agent (22). In apparent contradiction, blocking myosin II function via blebbistatin also leads to junction breakdown (23). Interestingly, inhibiting ROCK (kinase activated by RhoA) with Y-27632 is additionally detrimental to junction integrity (23, 24). As mechanical tension has been shown to directly increase the size of cell-cell adhesions (25, 26), RhoA signaling and actomyosin contractility evidently have a non-linear influence on the behavior of AJs and barrier function. Another member of the Rho GTPase family, Rac1, is also frequently implicated in the regulation of vascular function and endothelium homeostasis. Rac1 activates the WAVE regulatory complex to drive Arp2/3-associated polymerization of branched actin networks at the cell boundary. Abu Taha *et al*. (2014) observed that Rac1 induction facilitates the bridging of intercellular gaps and formation of stable junction contacts (1). Further, junction integrity cannot be maintained following blockage of Arp2/3 (23). By inducing junction tearing, Rajput et al. (2013) demonstrated that with decreasing Rac1 expression, gap closure was increasingly impaired (27). Conversely, significant upregulation of the Rac1 pathway via platelet-activating factor apparently causes widespread barrier disruption and cadherin endocytosis (11, 28).

The effects of both RhoA and Rac1 signaling are seemingly contradictory across the literature, promoting both junction strengthening and breakdown. In this work, we explore the formation, stability, and failure of endothelial contacts. We first propose a one-dimensional theoretical model to describe the three-way chemo-mechanical feedback between (i) VE-cadherin bond turnover, (ii) RhoA-mediated actomyosin contractility, and (iii) Rac1-drived actin polymerization, to uncover how endothelial gaps dynamically form and recover. We expand this framework to the analysis of a two-dimensional monolayer, resolving the contradictory influence of RhoA and Rac1 expression on junction stability. The model predicts how pharmacological modulation of signaling and downstream actin polymerization and cell contractility impacts junction stability, which we then validate experimentally by treating human umbilical vein endothelial cells (HUVECs) with thrombin, Y-27632, CK-666, and PAF. Ultimately, we demonstrate how a balance and crosstalk between RhoA and Rac1 signaling is essential in the regulation of homeostatic function.

## Results

### Two-way feedback between actomyosin contractility and VE-cadherin bond turnover governs cell tension and adhesion strength

The maintenance of vascular endothelial cell-cell junctions is predominantly governed by localization of VE-cadherin which form homotypic bonds with corresponding proteins on the membrane of neighboring cells (Figure 1A). Molecular-level studies indicate that cadherin bonds exhibit catch-bond behavior at low forces and slip-bond behavior at large forces (29–31). We may consider the force within a cadherin bond to be given by *F*_*b*_=*k*_*b*_*Δ*_*b*_, where *k*_*b*_ *is* the effective cadherin stiffness and *Δ*_*b*_ its change in length. Following from the recently developed catch-slip model of Novikova and Storm (2013), the dissociation rate of a single bond can be described by

**Figure 1.**
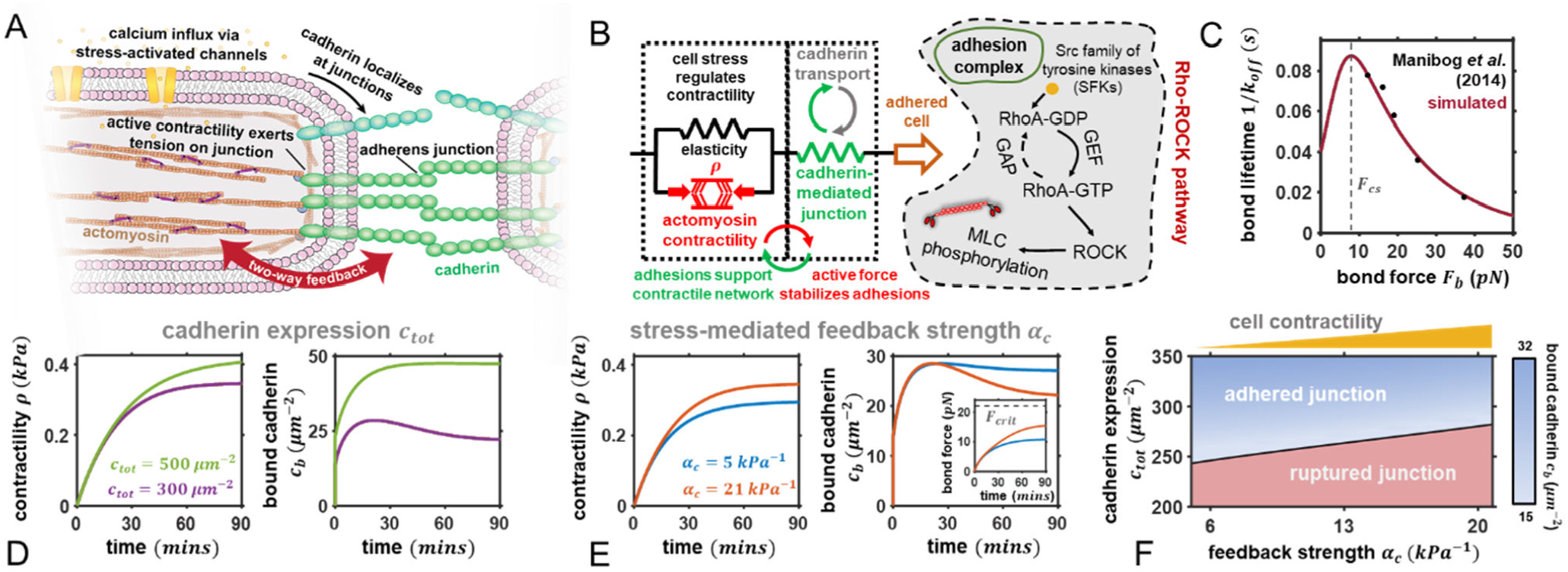
Feedback between cadherin activity and contractility in connected cells. (A) Schematic highlighting the feedback between actomyosin contractility and bound cadherin. In connected cells active cell tension stabilizes cadherin bonds which in turn anchor the actomyosin network and mediate signaling. (B) The mechanical model with actomyosin contractility, elastic stress, and cadherin-dependent junction stiffness. Biochemical Rho-ROCK pathway also shown whereby adhesion force downstream upregulates myosin phosphorylation; C) Experimental and predicted lifetime of a single VE-cadherin bond as a function of bond force. Cadherins present catch behavior under forces smaller than the catch-slip transition force *F*_*cs*_, while they show slip behavior at forces larger than *F*_*cs*_; Predicted dependence of contractility and bound cadherin density on D) cadherin expression *c*_*m*_ and E) adhesion-mediated feedback strength *α*_*c*_. For all simulations cells are initially in contact and the reference state (*t*=0 *) is ass*umed to be stress-free (*ρ* (*t*=0)=0) with no bonds yet formed (*c*_*b*_=*0*) and parameters *ρ*_*0*_=250, *Pa,c*_*tot*_*=*300*μm*^*−2*^ *an*d *α*_*c*_=21*kPa*−1 unless otherwise stated; F) Phase diagram of junction rupture/adhesion potential as a function of cadherin expression and feedback strength. The blue colormap shows the predicted steady state concentration of bound cadherin in connected junctions.

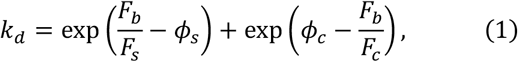

where *F*_*c*_ is the *‘*catch’ force, *F*_*s*_ is the *‘*slip’ force, and *ϕ*_*c*_ and *ϕ*_*s*_ are associated dimensionless catch-slip parameters. This model provides excellent agreement with experimentally measured lifetimes of single VE-cadherin bonds (30) as shown in Figure 1C, and naturally yields a catch-slip transition force (*F*_*cs*_), at which the bond lifetime reaches a maximum. When *F*_*b*_ *< F*_*cs*_ the lifetime increases as the force increases, characteristic of ‘catch’ behavior; when *F*_*b*_ *> F*_*cs*_ *th*e lifetime decreases as the force increases, also referred to as the ‘slip’ regime. The binding probability *P*_*b*_ for a VE-cadherin may then be expressed by *dP*_*b*_*/dt*=(1 *− P*_*b*_*)k*_*a*_ − *P*_*b*_*k*_*d*_, where the association rate *k*_*a*_ is assumed to be constant. Considering a junction with a sufficient VE-cadherin density *c*_*tot*_ and assuming uniform load distribution at a junction site (32), the turnover in the density of bound cadherin *c*_*b*_ can then be determined through a mean-field expression such that *c*_*b*_=*P*_*b*_*c*_*tot*_, whereby

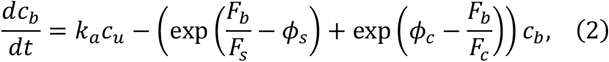

where *c*_*u*_ is the *u*nbound cadherin density and *c*_*tot*_=*c*_*b*_ *+ c*_*u*_. Further, we note a threshold force *F*_*crit*_ for unbinding beyond which bonds will rupture, reported to be on the order of 22 *pN* (30). Unbound cadherin may also be transported to and from the junction through passive and active processes (e.g. endocytosis), further explored in SI Section S1.

Adherens junctions connect directly to force generating actomyosin networks in the cell cytoplasm which can apply tension to the cadherin bonds and increase their stability. In turn, the forces generated at intercellular junctions trigger a range of biochemical processes that activate the Src family of kinases (SFK) as highlighted in Figure 1B. SFKs act on Rho-GTPases by controlling the activity of guanine nucleotide exchange factors (GEFs) and GTPase activating proteins (GAPs). Increased RhoA promotes Rho-associated kinase (ROCK), increasing myosin activity via phosphorylation of the myosin light chain and thereby promoting higher cell tension via actomyosin contractility. We have previously developed a chemo-mechanical feedback model to describe the interdependence of endothelial cell stress *σ*_*ec*_, signaling, and myosin-dependent force generation *ρ (33)*, extended here to consider explicit interactions with cell-cell junctions such that

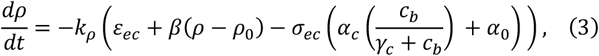

where *ε*_*ec*_ is the cell strain, *ρ*_0_ is a relative myosin motor density in the quiescent state, *k*_*ρ*_ is a kinetic constant governing the rate of myosin recruitment, and *β* denotes a chemomechanical coupling parameter regulating motor engagement. The *β*(*ρ - ρ*_0_) term ensures that the motor density in the absence of cell stress *σ*_*ec*_ is the quiescent value. *α*_*c*_ relates stress to signaling mediated by cadherin junctions (SFK, RhoA) where *γ*_*c*_ denotes the cadherin concentration for half-strength signaling, and *α*_0_ relates to junction-independent pathways such as calcium influx through mechanosensitive channels, which downstream activates myosin light chain kinase (MLCK) and promotes crossbridge cycling (34). Thus, the 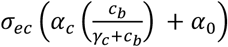 term promotes the recruitment of motors as governed by junction signaling and cell stress. We note the criterion: 0 < (*α*_*c*_ *+ α*_0_)*/β < 1*. Cell stress depends on both active and passive constituents, such that *σ*_*ec*_=*ρ + K*_*ec*_*ε*_*ec*_ where *K*_*ec*_ is the effective passive cytoskeletal stiffness. Initially, considering a cellular length of interest *L*_*ec*_ and assuming the reference state to be a stress-free connected junction, mechanical equilibrium reveals the following expression for the total system stress (for a detailed derivation see SI Section S2):

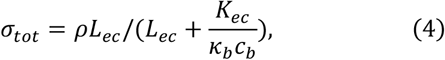

such that the force in an individual cadherin bond is then given by *F*_*b*_=*σ*_*tot*_*/c*_*b*_. Model parameters are provided in SI Section S4. We proceed to explore the interactions between adhesion stability and actomyosin contractility.

Our model predicts that as cell contractility increases, the density of bound cadherin *c*_*b*_ initially increases due to catch-type behavior stabilizing the bonds (Figure 1D). In turn, adhesion signaling feedback increases (via Eqn 4) which promotes higher contractility through myosin phosphorylation and cross-bridge cycling. However, with low cadherin expression (*c*_*tot*_=300 *µm*^*2*^), excessive tension drives the cadherin bonds into a slip-regime, increasing dissociation and weakening the adhesion. Increasing cadherin expression facilitates higher bond formation, and, although contractility is in turn predicted to increase through an upregulation in signaling, the cadherin bonds remain stable due to distribution of the load across more bonds. In agreement with our model predictions, promoting cadherin endocytosis via VEGF treatment has been shown to directly impair the stability of cell junctions (12). Reducing the maximum feedback strength *α*_*c*_ causes a reduction in contractility (Figure 1E), which can strengthen an adhesion by driving bonds back into a catch-regime. Under high tension if the bond forces exceed the threshold value *F*_*crit*_, complete adhesion rupture can occur. Thus, a balance between cadherin expression and signaling feedback is required to maintain connected junctions (Figure 1F): high cadherin expression coupled with actomyosin tension is associated with adhesion stability, while reductions in cadherin expression and excessive contractility can drive bonds into a slip-regime and, ultimately, failure. In support of these findings, thrombin, an activator of the RhoA signaling pathway, is a known effector of gap formation and junction permeability (20, 35). The opposing AJ-stabilizing influence of cell contractility predicted by our model has, until recently, not been widely appreciated. However, recent studies now indicate that tension induced by activated RhoA signaling can stabilize junctions, further reinforced through vinculin unfolding and protein recruitment (36); this latter pathway is explored in SI Section S1. In the next section, we proceed to identify how junctions can be restored in response to a rupture event.

### A critical level of Rac1-mediated actin polymerization is required to restore junction integrity post-rupture

In the event of junction failure, neighboring cells can reconnect by developing micro-protrusions, mediated by actin polymerization at the cell periphery (Figure 2A). Another member of the Rho GTPase family, Rac1, activates the WAVE regulatory complex to drive Arp2/3-associated polymerization of branched actin networks (Figure 2B) (37). Compressive stress *σ*_*P*_ generated by such polymerization can drive protrusion of the cell membrane. As such, the effectors Rac1 and RhoA work in opposition, governing protrusion and contraction, respectively. Interestingly, Rac1 is also suppressed by RhoA activity; signaling through ROCK activates Rac1 GAPs, and myosin activation locally prevents recruitment of Rac1 GEFs (38). We therefore propose a model to describe the level of branched actin network polymerization P such that

**Figure 2.**
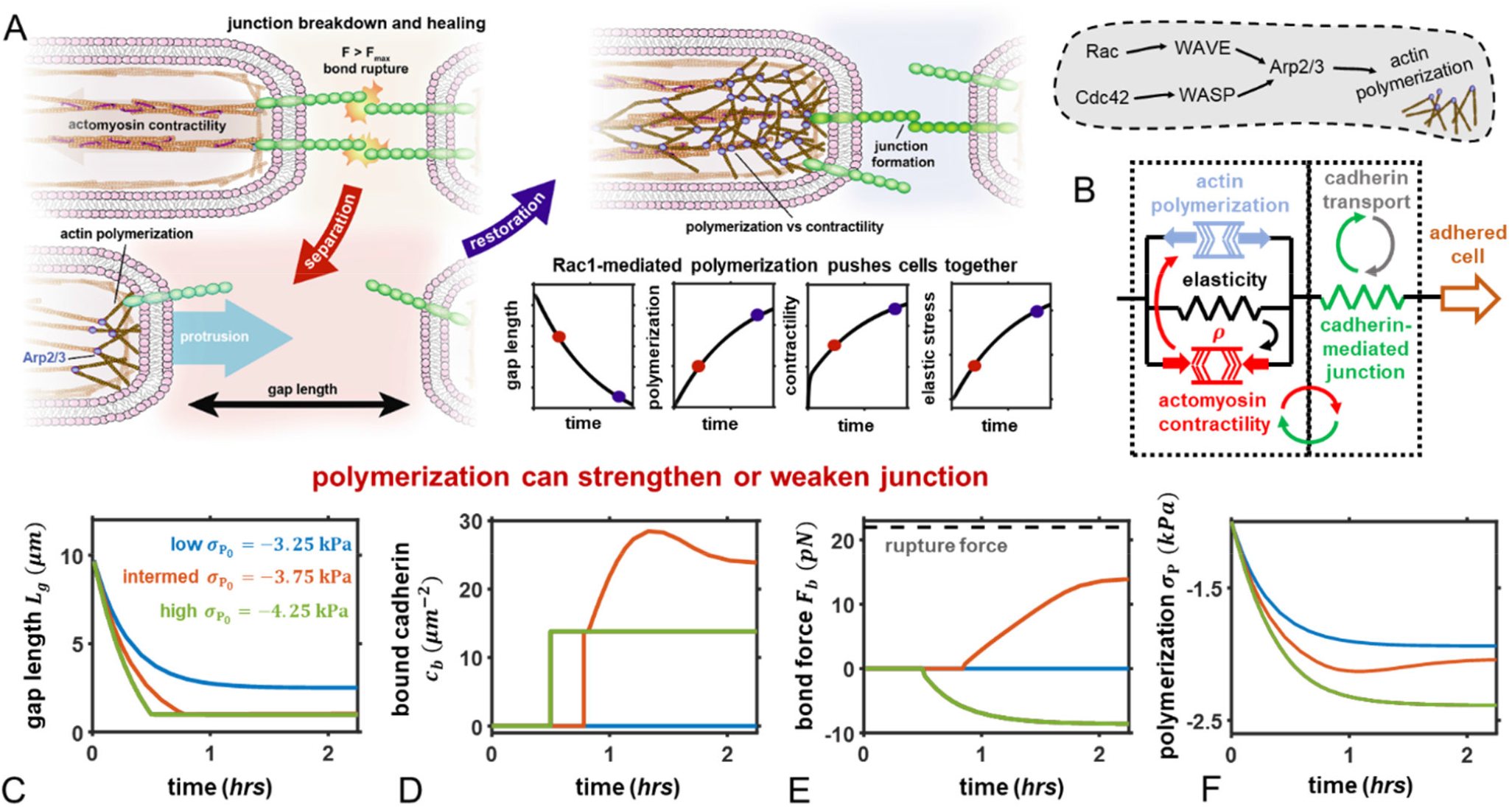
Actin polymerization restores ruptured junction. (A) Following adhesion rupture Arp2/3 mediated actin polymerization generates protrusive (negative by convention) stresses (red circle on plots) which can ultimately lead to junction restoration (blue circle on plots); (B) The mechanical model considers stresses induced by polymerization which are opposed by active and passive cell stresses. These parallel components act in series with the cadherin bonds which remodel in response to tension. Biochemical pathway for polymerization also shown whereby Arp2/3 is upregulated downstream of Rac and Cdc42; (C-F) A low polymerization-induced stress is predicted to be insufficient to reconnect cells and over-polymerization to prevent actomyosin contractility from stabilizing cadherin bonds: associated C) gap length, D) bound cadherin concentration, E) individual bond force, and F) polymerization-induced stress *μ*_P_. For all simulations cells are considered to be post-rupture and initially separated with a gap distance *L*_*g*_=10*μ m*. The reference state (*t*=0*) is ass*umed to be stress-free (*ρ* (*t*=0) = − *σ*_P_ (*t*=0)=1*kpa*) with parameters *ρ* _0_=1*kpa, c*_*tot=*_ 300*μ m*^*−2*^ *an*d *α*_*c*_=20 *kpa*^*−1*^ unless otherwise stated.

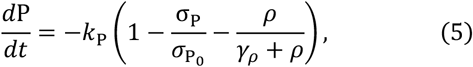

where 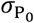 is the maximum potential compressive stress induced by actin polymerization, associated with a maximum level of Rac1 signaling. The polymerization-induced stress 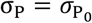, and *k*_P_ is a kinetic constant. The first bracketed term describes the self-inhibition of Rac1 (and downstream polymerization) at high expression levels (39) (Figure S3A), and the second term describes the antagonistic interactions between RhoA and Rac1 as determined by suppression constant *γ*_*ρ*_; with increasing contractility *ρ* (proxy for RhoA expression), Rac1-mediated actin polymerization will reduce (Figure S3B). In an open system (unconnected cells), stress equilibrium mandates that the cell strain is given by *ε*_*ec*_=−(*ρ +* σ_*P*_*)/k*_*ec*_. Therefore, actin polymerization and actomyosin tension directly compete to enable cell protrusion or contraction (Figure 2A) such that the junction can be restored if an inter-cell gap of reference length *L*_*g*_ is sufficiently reduced (i.e. *L*_*ec*_*ε*_*ec*_ *≥ L*_*g*_). Once the cells come into contact, bond formation and reinforcement can occur (1), with the expression for stress equilibrium then given by:

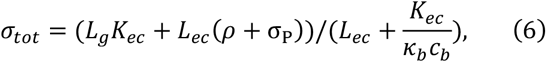

with a detailed derivation provided in SI Section S2. Again, the junction will rupture if the bond force *F*_*b*_=*σ*_*tot*_*/c*_*b*_ exceeds a critical value *F*_*crit*_, returning to an open system. Following junction rupture, our model predicts that a critical level of actin polymerization is required to successfully close the gap (Figure 2C). If polymerization is too low, the gap distance cannot be sufficiently reduced and adhesions cannot form (Figure 2D,E), supported by our previous findings (23) in which inhibition of actin polymerization limited junction recovery. Although contractility also remains low (Figure S3C) due to lack of SFK signaling, the cortical tension mediated by *α*_0_ and cytoskeletal elasticity are sufficient to counter the protrusive forces from polymerization (Figure 2F). If the polymerization-induced stress becomes dominant and the gap closes, cadherin bonds are then predicted to form (Figure 2D) and the feedback between contractility and adhesion reinforcement can evolve. As contractility increases with increased RhoA expression from junction signaling, polymerization reduces due to Rac1 suppression (via Eqn 5); our model also indicates that junction compression from over-polymerization may prevent the transmission of tensile forces through cadherin bonds and therefore limit adhesion stability, reinforcement, and formation of actomyosin networks (Figure 2C-F). In support of this prediction, Knezevic et al. (2009) report that platelet-activating factor, an activator of the Rac1 pathway, drives junction breakdown as characterized by cadherin endocytosis (28). Next, we explore how dynamic junction rupture and repair can emerge as dependent on interactions between RhoA and Rac1 signaling and feedback.

### Dynamic gap behavior and junction stability emerge at different levels of RhoA and Rac1 expression

Having indicated how RhoA-driven contractility and Rac1-mediated actin polymerization interact to govern endothelial junction strength and recovery, we proceed to investigate how the junction state depends on crosstalk between these processes. Our model predicts that dynamic behavior emerges within a critical feedback strength and polymerization stress range (Figure 3A); protrusion facilitates cell-cell contact, enabling spontaneous cadherin bond formation. As the density of bonds increases, the stress-mediated signaling feedback strength also increases which upregulates signaling from SFKs and causes downstream myosin motor phosphorylation and cross-bridge cycling (Figure S4B). In agreement, using high-speed imaging Yamada and Nelson (2007) observed that Arp2/3-induced lamellipodia rapidly give way to stress-activated RhoA recruitment following intercellular contact, driving a strengthening of nascent cell contacts (40). With higher contractility within a catch regime, bond dissociation is reduced which overall promotes a higher bond density. In conjunction, the polymerization stress is reduced via RhoA-associated Rac1 suppression (Figure S4A). However, development of further contractility then pushes the adhesion into a slip-regime, driving rapid bond dissociation which in turn increases the force acting on individual bonds (less distributed). Rupture ensues when this force exceeds the threshold value, causing cell-cell separation. A loss of adhesion-mediated signaling then lowers contractility (Figure S4B) in turn facilitating a restoration of the polymerization-induced stress and allowing the cells to reconnect. This process cycles repeatedly over a calculable timescale that depends on the level of RhoA and Rac1 signaling (via *α*_*c*_ and 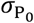, respectively), closely resembling the ‘stick-slip’ cycle often reported in mesenchymal-type migration (41, 42). Exploring the phase space of these two parameters facilitates a mapping of the junction state (Figure 3B). A high level of Rac1 signaling, coupled with low signaling feedback from adhesion stretching, is predicted to mediate junction weakening in the form of a closed cell-cell gap but an absence of bond tension and stability. With increasing feedback *α*_*c*_, the junction strength increases due to higher levels of actomyosin force generation acting on the adhesions, leading to bond stabilization and reinforcement (via Eqn 2,3) in a catch-regime. Impairment of Arp2/3-mediated polymerization is predicted to prevent gap closure, as insufficient stresses are generated to drive development of protrusions (43). Moreover, as noted there is a critical balance of Rac1 and RhoA signaling that our model suggests will lead to the emergence of dynamic behavior.

**Figure 3.**
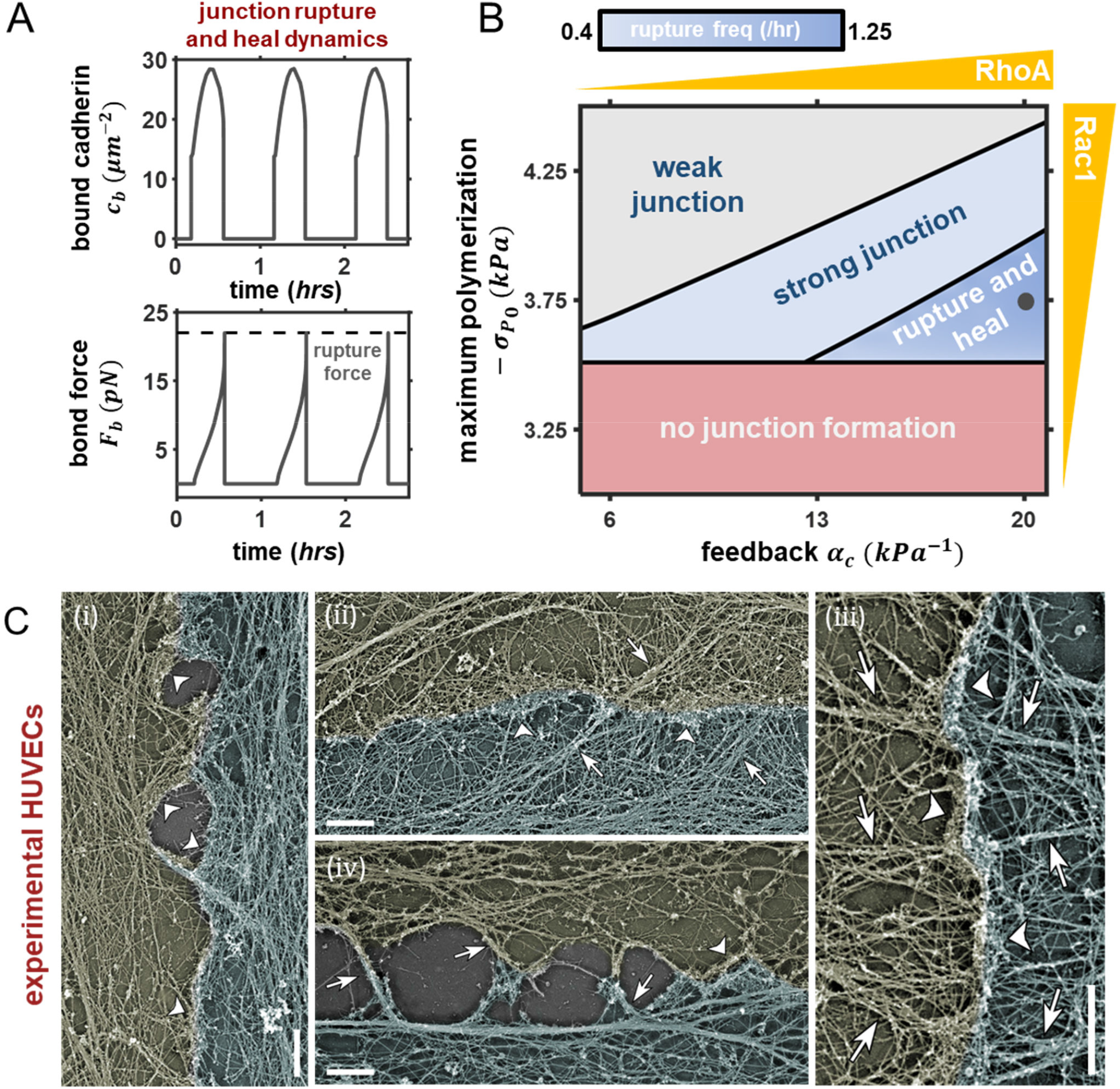
Phase diagram for junction stability. A) Transients in bound cadherin density and individual bond forces during dynamic activity (*α*_*c*_ =20 *kpa*^*−1*^ = 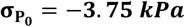; B) Phase diagram of junction behavior as driven by actin polymerization and adhesion-mediated feedback. The grey marker indicates the parameters for (A); C) PREM images showing cytoskeleton organization associated with (i) perforated, (ii/iii) connected, and (iv) tearing junctions. Arrows and arrowheads mark actin bundles and branch networks, respectively. Scale bars, 1*µm*. C-iii is reproduced with permission from our prior work (23) under identical conditions. Contacting cells are lightly pseudo-colored to highlight cell boundary, which was identified based on immunogold VE-cadherin labeling.

To assess our model predictions for the dynamic rupture and recovery of endothelial junctions, we performed platinum-replica electron microscopy (PREM), in which monolayers of cultured human umbilical vein endothelial cells (HUVECs) were extracted, fixed with glutaraldehyde and coated with a thin layer of platinum before imaging with transmission electron microscope (see Methods). This technique permits high resolution imaging of cytoskeletal network organization (44, 45). Under control conditions, we observed that the organization of the cytoskeleton was indicative of the junction state (Figure 3C) at different stages of the rupture/heal cycle. Continuous cohesion of two neighboring cells is characterized by the presence of branched actin networks at the cell-cell junction and actin filament bundles (perpendicular, oblique or parallel) in the junction vicinity (Figure 3C-ii,iii). Apparent post-rupture junctions are mainly associated with actin bundles that often form intercellular bridges (Figure 3C-iv), whereas intercellular gaps that appear to be in a process of gap closure are often outlined by dense branched actin networks (Figure 3C-i). In previous work we also identified a high density of Arp2/3 in regions containing branched actin networks supporting their protrusive nature (23). These data correspond to our model predictions whereby sufficient polymerization is required to close gaps (Figure 3C-i) and maintain junctions, such as observed near the connected cell boundary in Figure 3C-ii. As a junction develops, our model indicates that adhesion-mediated signaling drives increased actomyosin activity, which in turn applies tension to the cadherin bonds and promotes stability. This behavior is observed in Figures 3C-ii and 3C-iii, where oblique or perpendicular contractile actomyosin fibers, respectively, are connected to the cell-cell boundary. As the feedback evolves, thick stress-fibers can form which impose high tension and may tear the junction apart (Figure C-iv). Again, this agrees with model predictions whereby the evolution of high actomyosin stress drives cadherin bond dissociation and junction failure (Figure 3A). Our experimental images also clearly reveal that the junction state varies spatially along the boundary. Motivated by these observations, in the next section we proceed to extend our computational model to the analysis of a 2D monolayer exhibiting dynamic non-uniform active behavior.

### Rupture initiates at multi-cell junctions in an endothelial monolayer with healing mediated by recovery of actin polymerization

We proceed next to extend our framework to the analysis of junction dynamics in an endothelial network. Prior, we have implicitly assumed that polymerization, contractility, and adhesion activity are uniform along a protruding cell boundary. However, due to geometric constraint and fluctuating cell activity, variance will emerge when considering the behavior of an endothelial monolayer. To address such non-uniform dynamics, we develop a non-linear 2D spring lattice model as motivated by our previous work (46). We discretely consider (i) independent protrusions comprised of active cytoskeletal (polymerizing/contractile) elements (Figure 4), (ii) adherens junction elements that can rupture, remodel, and reform, and (iii) passive membrane/cortical actin elements that connect the protrusions. Elements in the lattice are connected by nodes, and associated chemo-mechanical behavior is governed by our proposed model (Eqns 1-5). The system is solved using a Newton-Raphson iterative scheme (see Methods) such that mechanical equilibrium is achieved at every node at every time point in a quasi-static analysis. We can then further extend to assemble a representative volume element of a symmetric cell within the monolayer (Figure 4; Figure S5). Model parameters are provided in Table S2.

**Figure 4.**
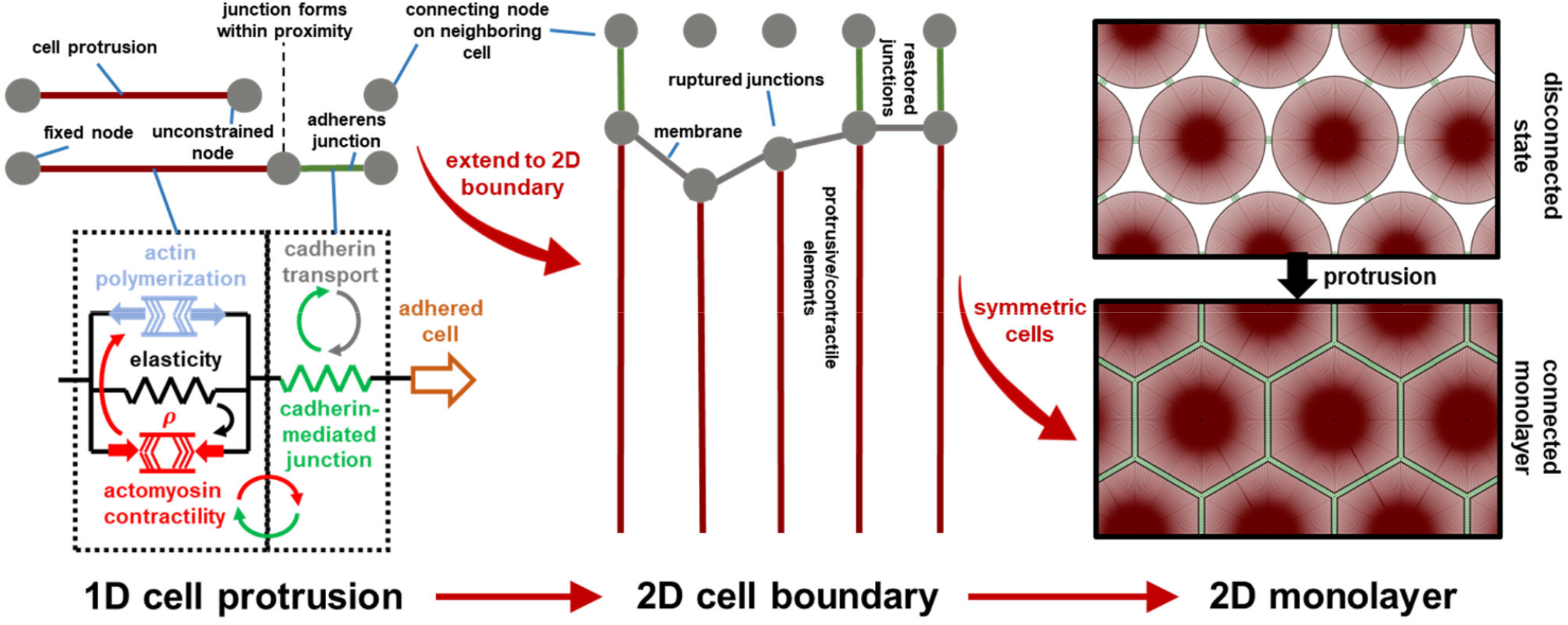
Schematic of 1D model extension to a 2D lattice-based monolayer. The 2D model discretely considers independent protrusions comprised of cytoskeletal (polymerizing/ contractile) elements (red), adherens junction elements that can rupture and reform (green), and membrane/cortical actin elements that connect the protrusions (grey). Elements in the lattice are connected by nodes and the system is solved using a Newton-Raphson iterative scheme. Cells protrude in response to actin polymerization to close gaps.

We cultured a confluent monolayer of HUVECs on collagen gels and studied the evolution of cell-cell junctions (see Methods). We observed that gaps form at endothelial junctions, grow in size, and heal over time (Figure 5A). In agreement, our simulations of monolayer behavior suggest that when a gap forms, separation propagates along the boundary which further increases the gap size (Figure 5B). Then, due to the loss in contractility (associated with adhesion rupture and signaling downregulation) and increasing polymerization, the cell membrane protrudes to reestablish contact and bonding. Experimentally, we found that junctions are more frequently disrupted at vertices (multi-cellular junctions) than at borders (bicellular junctions), with quantitative measurements demonstrating that the probability of observing a gap at a vertex is approximately nine times higher than that at borders (Figure 5C). Our model indicates that the forces exerted on cadherin bonds are significantly higher at vertices than at two-cell borders (Figure 5D), which can drive bond rupture and junction failure; this emerges from a combination of constraint, alignment of the cortex and associated stress, and the high regional stretch required to develop contact at the vertex. To further assess the development of such vertex stress concentrations, we developed an analytical model which reveals that a stress singularity can arise at the multi-cellular junction when there is a mismatch between the cytoskeletal and adhesion stiffness. This analysis is discussed in detail within SI Section S3. Gaps were observed to form at the HUVEC vertices approximately once per hour, with ruptures persisting over 30 minutes (Figure 5E). Our simulated timescales provide excellent agreement with these data, further supporting the validity of our framework.

**Figure 5.**
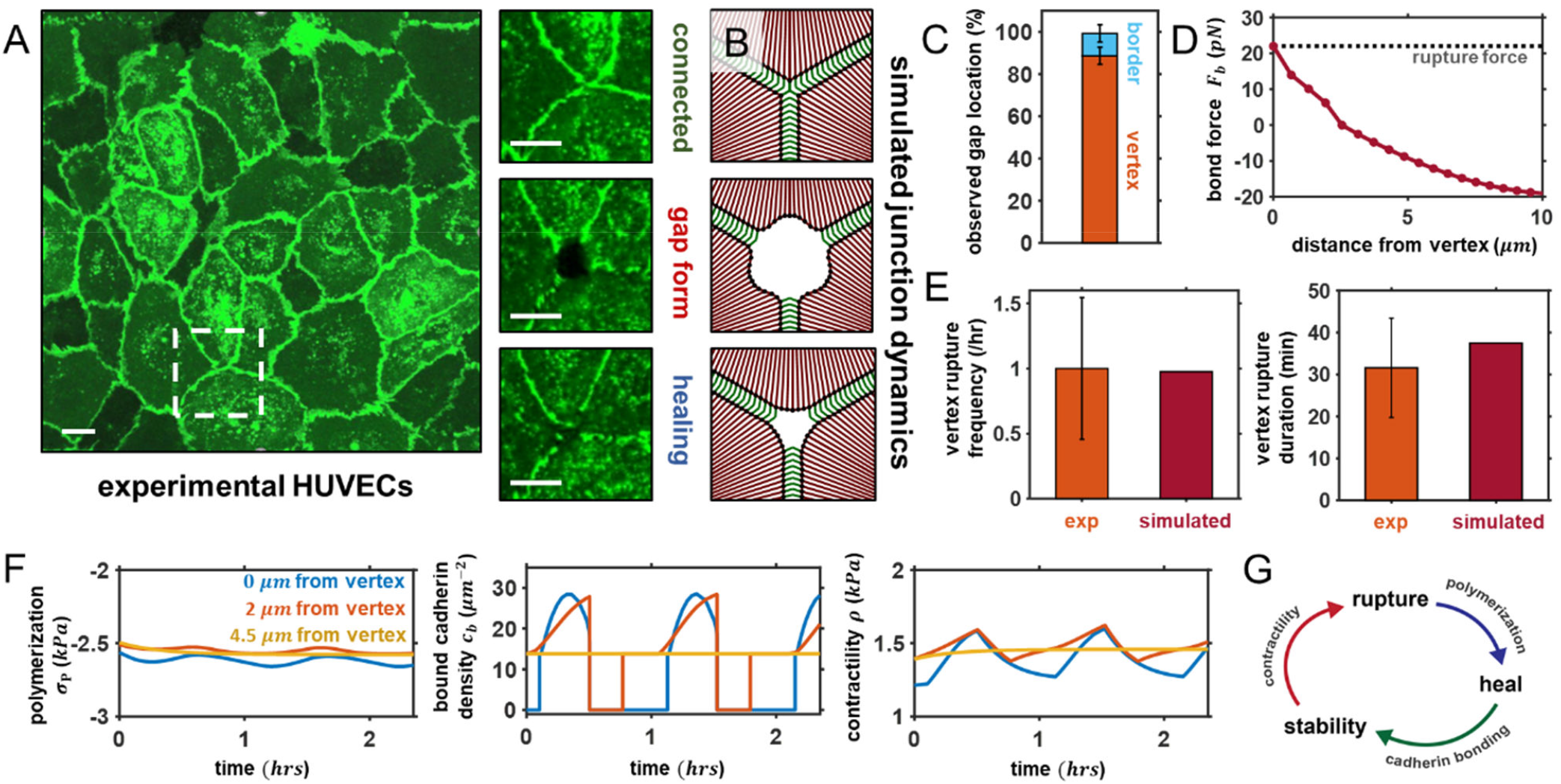
Simulations of Cell Network Dynamics. A) Dynamic behavior of HUVECs expressing VE-cadherin-GFP cultured on a thin collagen gel, showing typical formation and healing of gaps at a multi-cell vertex. Subregion frames recorded at 0 min, 70 min, and 130 min (top to bottom). Scale bar: *µm*; B) Sample gap formation and healing within the simulated monolayer; C) Experimentally observed gap locations in the HUVEC monolayer with rupture predominantly occurring at multi-cell vertices; D) Predicted bond force along the adhered cell edge prior to rupture event indicating highest force is localized at the vertex; E) Experimental and simulated rupture frequency at a vertex and duration of rupture; F) Predicted polymerization, bound cadherin density, and contractility along the cell edge; G) Schematic of rupture and recovery cycle as critically dependent on contractility, polymerization, and cadherin bond stability. For all simulations *α*_*c*_= 15 *kPa*^−1^ and 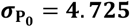 unless otherwise stated.

Our analysis clearly indicates that cell activity is non-uniform along boundaries within a monolayer. Growing tension at vertex bonds causes them to enter a slip regime and subsequently the local adhesion fails (Figure 5F). As the cell locally retracts in this region, additional stress is imposed on neighboring adhesions which causes the separation to propagate along the boundary. The rupture is arrested when the critical bond force within an adhesion is not surpassed and contact can thus be sustained. Over time, the adjacent membrane is predicted to protrude due to actin polymerization and loss of contractility which enables cell contact and adhesion until the whole boundary is restored. With junctions restored along the cell surface, adhesions strengthen and contractility increases until rupture occurs again (Figure 5G).

### Pharmacological treatments modulating cell contractility and actin polymerization drive weakening and failure of junctions

We further investigate the role of Rac1 and RhoA signaling (implicitly considered through 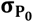 and *α*_*c*_, respectively) on the maximum gap radius that emerges in the monolayer (Figure 6A); simulations suggest that an increase in Arp2/3-mediated polymerization can reduce gap formation and limit rupture propagation, while high signaling feedback to myosin phosphorylation and contractility drives larger gaps and further separation. In agreement with our findings, angiopoietin, a Rac1 activator and RhoA inhibitor (47), has been shown to significantly reduce both endothelial gap size and rupture propagation (48). We can relate specific levels of RhoA and Rac1 expression to cell behavior in response to associated pharmacological treatments. At a critical balance between these signals, stable junctions with a high bound cadherin density are maintained at steady state (Figure 6B-D). Our model suggests that increased activation of RhoA and downstream actomyosin contractility will drive increased gap formation and junction breakdown. In support of these findings, we found that treatment with thrombin, an activator of the RhoA signaling pathway, increases junction permeability (Figure 6C). Conversely, a reduction in cell contractility is predicted to reduce the cadherin bond density and thus weaken cell-cell adhesion. To test this prediction, we treated HUVECs with ROCK inhibitor Y-27632, which prevents downstream activation of non-muscle myosin II.

**Figure 6.**
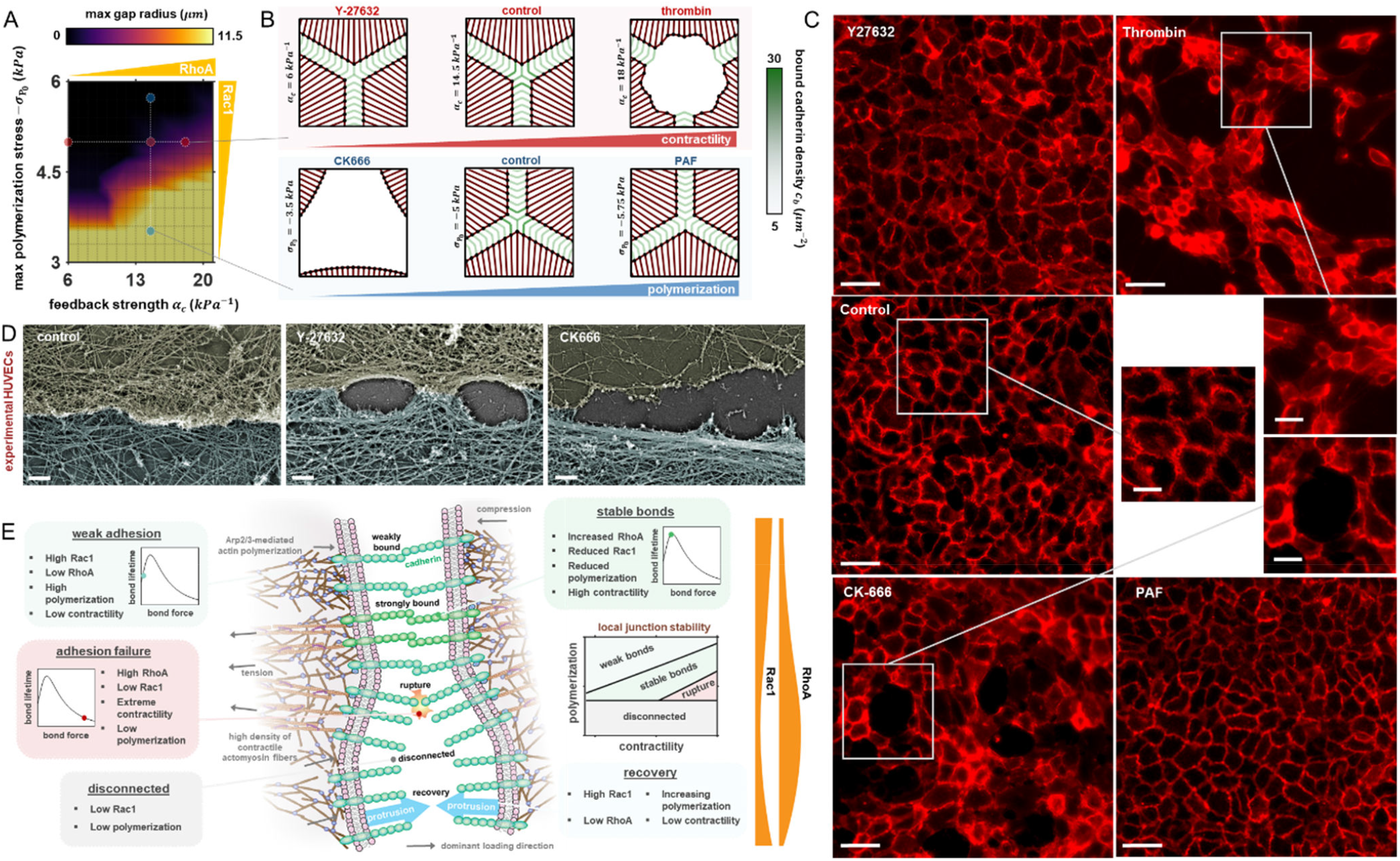
Junction state is governed by RhoA and Rac1 signaling. A) Maximum predicted gap radius as a function of feedback strength *α*_*c*_ and maximum polymerization-induced stress 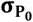; B) Predicted influence of pharmacological treatments on junction stability and gap formation. Scale bar, 25*µm*; C) Organization of HUVEC monolayer in response to pharmacological treatment with Y-27632, thrombin, control (untreated), CK-666, or PAF. Scale bar, 50*µm*. Red fluorescence shows CD31. Inset scale bar, 25*µm*; D) PREM images of HUVECs showing cytoskeleton organization and junction state associated with control conditions and treatment with Y-27632 or CK666. Contacting cells are pseudo-colored to highlight cell boundary, which was identified based on immunogold VE-cadherin labeling. Scale bars, 0.5*µm*; E) A balance of RhoA and Rac1 signaling is essential to the maintenance of endothelial junctions with variations driving junction weakening or failure.

This led to the emergence of non-adhered regions despite a high density of polymerized actin (Figure 6D), in agreement with our model’s indication of weakened adhesion. Walsh *et al*. (2001) also demonstrated that limiting ROCK activation is detrimental to junction integrity (24). RhoA and Rac1 have opposing effects on cell protrusion, and as such simulations indicate that upregulation of Rac1 (e.g. through endothelial cell treatment with platelet-activating factor (PAF)), also causes junction weakening due to a reduction in VE-cadherin bond tension (Figure 6B). In agreement, PAF has been reported to cause widespread barrier disruption and cadherin endocytosis (28). Such endocytosis is reported to occur under low junction tension (49), additionally explored in SI Section S1. Our model also suggests that insufficient actin polymerization can inhibit junction formation, as cells cannot sufficiently bridge the gap between their neighbors. In support of this finding, we show that large voids emerge following blockage of Arp2/3 (23) and associated actin branching through treatment with CK666 (Figure 6C,D). Abu Taha *et al*. (2014) also report that Rac1 mediates the bridging of intercellular gaps and formation of stable junction contacts (1). In summary, our model indicates that both RhoA and Rac1 expression have a bimodal influence on junction integrity which should be appreciated when drawing conclusions from pharmacological regulation. Further, a balance between these signaling processes is required for the maintenance of stable adhesion (Figure 6E).

## Discussion

In this study, we propose a chemo-mechanical model to investigate how endothelial gaps are regulated within the microvasculature. Junction breakdown and recovery depend on the three-way feedback between active force generation, cadherin bond turnover, and actin polymerization. Assembly and disassembly of adherens junctions (AJs) at the cell-cell interface is force-dependent, with cadherin proteins exhibiting catch-bond behavior at low forces and slip-bond behavior at high forces (30). In connected cells, actomyosin contractility is therefore shown to significantly affect adhesion stability, such that actively generated tension can increase bond lifetime but excessive stress causes bond dissociation and adhesion rupture (Figure 6E). The recruitment of myosin motors is a mechanosensitive process that occurs downstream of signaling from AJ tension. Thus, a feedback loop emerges between adhesion stability and actomyosin contractility, whereby stretching of cadherin bonds upregulates RhoA signaling (via coupling parameter *α*_*c*_) to promote myosin-mediated force generation, in turn transmitting higher tension to the junction.

Following the formation of endothelial gaps, Rac1-associated actin polymerization can aid junction recovery by protruding the membrane to facilitate cell-cell contact; this process is further advanced through the reduction in cell contractility associated with loss of adherens junctions and related signaling. Our model indicates that insufficient Rac1 signaling (implicitly considered through 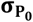) may prevent gap closure in agreement with our previous findings (23), while high Rac1 expression can lead to weak junctions characterized by a low bound cadherin density. The work of Knezevic *et al*. (28) supports this latter result, who observed that high activation of Rac1 drives junction weakening. Our simulations suggest that a balance between RhoA and Rac1 signaling is required to maintain a stable junction state (Figure 6B); reduced RhoA or Rac1 expression led to weak adhesions or perforated cell-cell contacts. These results were validated through platinum-replica electron microscopy (PREM) on cultured HUVEC monolayers, a technique which facilitates high resolution imaging of cytoskeletal networks. We observed that perforated and weak junctions were associated with regions of low actin density and alignment (Figure 3C), while strong, stable junctions exhibited dense and aligned networks. Simulations also suggests that high levels of RhoA expression and associated actomyosin contractility drive adhesion failure; in agreement our PREM images show that thick stress-fiber bundles were assembled in regions that junctions had started to tear apart. The model proposed in this study can therefore help to resolve the seemingly contradictory effects reported for RhoA and Rac1 expression, by which through coupled interactions they can drive junction assembly, failure, recovery, or stability as dependent on their associated levels and crosstalk.

We next extended our framework to explore how spatial variations in gap formation can emerge along cell-cell boundaries within an endothelial monolayer. We cultured a confluent monolayer of HUVECS on collagen gels and observed the propagation of gaps along a boundary and subsequent healing in agreement with the timescales predicted by our model. Gaps had a tendency to form more frequently at multi-cellular junctions than at two-cell borders. Our numerical simulations suggest that the forces exerted on cadherin bonds are significantly higher at vertices than at bicellular boundaries, which can drive bond rupture and loss of adhesion, thereby promoting increased junction failure at multi-cellular junctions. Further, in SI Section S3 we show analytically that at a vertex, stress concentrations can arise from a mismatch between adhesion and cytoskeletal stiffness. The magnitude of the stress singularity at the triple-cell junction increases as the stiffness mismatch increases. Our numerical analysis also predicts the impact of pharmacological modulation of signaling and downstream cytoskeletal remodeling on junction stability, with larger gaps predicted to form with high RhoA expression or low levels of actin polymerization. We then experimentally validated predictions by treating HUVECs with thrombin, Y-27632, CK-666, and PAF (Figure 6C,D). Future studies could further test our model by exploring Src-specific inhibition and consequent changes to actomyosin contractility and gap formation (50).

Adherens junctions play a critical role in supporting many physiological processes, including collective cell migration (51) and morphogenesis (52). Clearly, such biological systems could be analyzed using our framework to provide mechanistic insight into their dependence on the crosstalk between active contractility and actin polymerization. Future advancements should also focus on detailing the feedback between integrin signaling and cadherin bond reinforcement (53), building on our previous work exploring the interdependence of focal adhesion development and cell force generation (54, 55). The role of cell-cell interactions (56) in endothelium homeostasis should also be investigated, in addition to region-specific loading in the vascular (57) and myocardial (58) endothelium. Further, in our main analysis we limit ourselves to the consideration of a constant total cadherin density at each junction site; however, the influence of active transport processes (endo- and exocytosis) on adhesion strength is explored in SI Section S1, with our analysis suggesting that force-mediated signaling can have a secondary indirect effect on bond stability by driving cadherin reinforcement through reduced endocytosis.

The activity of the Rho family of GTPases in the endothelium has major connotations for vascular behavior. Vascular barrier function can be compromised during inflammation, with the inflammatory mediator thrombin driving endothelial hyper-permeability through RhoA activation (20, 35), thereby promoting extravasation of blood constituents. Our analysis suggests that this occurs through increased actomyosin contractility pushing cadherin bonds into a slip regime and subsequent rupture. Previous experiments have also identified that Rac1 suppression alone is sufficient to disassemble endothelial junctions (59). Our simulations indicate that this effect is two-fold: reduced protrusion from a loss of actin polymerization lowers the compressive stress acting on the junction, thereby increasing bond tension and dissociation, and sufficient protrusive forces cannot then be generated to reconnect the cells. The formation of atherosclerotic lesions has been linked to endothelial dysfunction (60), with such dysfunction also dependent on the RhoA/ROCK pathway (61) and cell contractility. Conversely, pharmacological inhibition of Rac1 has been reported to rescue endothelial dysfunction (62). As plaque development initiates early in life (63), increased preventative measures should be supported to reduce the burden; future advancements could see our model extended to predict endothelial dysfunction and to develop a computational tool to advise on patient risk and early diagnosis of atherosclerosis. Beyond vascular disease, endothelial junction dynamics are also critical to cancer progression and metastasis. Tumor cells have been observed to exploit disrupted connectivity at multi-cellular junctions to transmigrate through the endothelium (64). Prior work has considered endothelial gap formation (46) by modeling constant contractile and protrusive forces that were randomly activated or deactivated over time, which provides an understanding of how junctions could break down to facilitate cancer cell extravasation. While the biochemical or biophysical signals that governed this process were unclear, there appeared to be a mechanism by which the tumor cells could identify vertices to increase their extravasation potential. The current work entails a significant advance over prior analyses, by integrating the dynamic remodeling of actomyosin networks and cell-cell adhesions with signaling feedback governed by cell mechanosensation. Simulations suggest that gap formation is critically dependent on stress-sensitive adhesion-mediated signaling that upregulates actomyosin contractility, with junction restoration driven by a subsequent recovery in actin polymerization. We further highlight a mechanism by which junctions are most likely to break down at multi-cellular interfaces; cell constraint drives higher actomyosin tension and the emergence of concentrated adhesion stress in these regions. In summary, our findings suggest that the three-way feedback between actomyosin contractility, VE-cadherin bond turnover, and actin polymerization governs the regulation of endothelial gaps. Our proposed model can help guide the development of therapeutics that target the Rho family of GTPases and downstream active mechanical processes.

## Methods

### Platinum-replica electron microscopy (PREM)

For PREM experiments, HUVECs (Lonza, CC-2519) were cultured in Endothelial Cell Basal Medium (Lonza, CC-3121) with supplements (Lonza, CC-4133) and maintained for no longer than six passages. For experiments, HUVECs were plated on coverslips coated with *∼*50 μg/ml (5 mg/cm^2^) collagen from rat tail (BD Biosciences, 354236). Y-27632 (Y100500; Toronto Research Chemicals) was prepared from 10-mM stock in water and cells treated were treated with 50 *μM Y-276*32 in culture medium for 1-hr. CK666 (SML0006; Sigma-Aldrich) was prepared from 10-mM stock in DMSO and used at a concentration of 100 *μ*M for 40*–6*0 min. Sample preparation for PREM was performed as described previously (45). In brief, HUVEC monolayers were extracted with 1% Triton X-100 in PEM buffer (100 mM PIPES-KOH, pH 6.9, 1 mM MgCl_2_, and 1 mM EGTA) containing 2% polyethelene glycol (mol. wt. 35,000), 5 µM phalloidin and 2 µM taxol for 3 min, washed with PEM, incubated for 30 min with a primary mouse monoclonal antibodies for cadherin-5 (VE-cadherin) (BD Biosciences, 610251) in the PEM buffer containing 5 µM unlabeled phalloidin and 2 µM taxol, washed with the PEM buffer and fixed with 0.2% glutaraldehyde in 0.1 M Na-cacodylate, pH 7.3. After quenching with 2 mg/ml NaBH_4_ in PBS for 10 min cells were blocked with 1% BSA in buffer A (20 mM Tris-HCl (pH 8), 0.5 M NaCl, 0.05% Tween 20), stained with secondary anti-mouse IgG antibodies conjugated to 12 or 18 nm colloidal gold (Jackson ImmunoResearch Laboratories) and postfixed with 2% glutaraldehyde in 0.1 M Na-cacodylate, pH 7.3. Fixed cells were sequentially treated with 0.1% tannic acid and 0.2% uranyl acetate in water, critical point dried, coated with platinum and carbon, and transferred onto electron microscopic grids for observation. PREM samples were analyzed using JEM 1011 transmission electron microscope (JEOL USA, Peabody, MA) operated at 100 kV. Images were captured by ORIUS 832.10W CCD camera (Gatan, Warrendale, PA) and presented in inverted contrast. Pseudocolors were applied using Hue/Saturation tool in Adobe Photoshop to avoid obscuring structural details.

### Tracking monolayer dynamics

To mimic endothelial monolayer in a physiological state and enable high-resolution confocal live-imaging of endothelial gaps, HUVEC monolayers were cultured on top of thin layers (∼60 µm) of collagen. Briefly, HUVECs (Lonza, expanded to passage 3) were transduced with GFP tagged VE-cadherin (65) and expanded and cryopreserved at passage 6. Transduced HUVECs (350,000 at passage 7) were seeded on top of collagen substrate (rat tail type I, Corning, 2.5 mg/ml, thickness of ∼60 µm) formed in the central region of Mattek glass-bottom dish prior to seeding. Two days following seeding, a uniform semi-permeable HUVEC monolayer with physiologically relevant permeability (tested with dextran diffusion) was formed. The dynamics of gap opening were captured over 3 hrs (every 10 min) using live confocal imaging (Olympus FV1000, 63X oil objective) to image the dynamics of VE-cadherin under physiological incubation condition (37°C, 5% CO2). To improve the quality of imaging, in a few cases Alexa Fluor 647-conjugated PECAM-1 antibody (Hu CD31, BD BioSciences) was applied to the monolayer and washed to image junctional dynamics for a short period of time (∼1 hr). To obtain enough measurements for statistical analysis, six regions on three separate dishes containing fully confluent HUVEC monolayer were experimentally tested. Fluorescence time-lapse images of cell junctions were manually analyzed using ImageJ to measure frequency and duration of gaps identified as void regions along junctions with sizes greater than ∼2 µm. For pharmacological perturbation, HUVECs were plated at 4,200 cells/mm2. After 24 hours, cells were treated with either thrombin1 (0.01U/mL; Millipore-Sigma, SKU: 10602400001), CK-6662 (200nM; EMD Millipore, 182515), PAF3 (100nM; Fisher Scientific Tocris, 29401), Y-276324 (30uM; EMD Millipore, 5092280001), or left untreated overnight. Next day, wells were washed once with DPBS, then incubated for 10 minutes with Alexa Fluor 647 Mouse Anti-Human CD31 (BD Bioscience, 561654) at 1:20 dilutions in EGM-2 media. Cells were rinsed twice with EGM-2 media, then imaged by fluorescent microscopy using a Nikon Eclipse Ti-2E microscope.

### Cell network dynamics model development

Cells are assumed to form repeating hexagonal shapes in a monolayer when spread and adhered (Figure S5). Taking advantage of the symmetry in this system, we model one-twelfth of an initially-circular cell body. The cell is discretized into nodes, connected by a system of elements that describe the chemo-mechanical behavior of the cytoskeleton, membrane/cortical actin, and cell-cell adhesions (Figure 4). Considering a given element *E*_*ij*_ connected to nodes ***n***_*i*_ and ***n***_*j*_, its length *L*_*Eij*_ may be determined by computing the current distance between the nodes, with *L*_*Eij*_=∥***n***_*i*_ *− **n***_*j*_∥. With a reference length 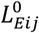, the force vector associated with element deformation may then be determined by 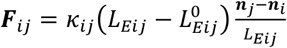, where *k*_*ij*_ is the spring stiffness of the element in units *N/μm*. Element forces act on the connected nodes, such that the total force vector for a given node ***n***_*i*_ is described by ***F***_*i*_=*∑*_*j∈neighbor*(*i*)_ ***F***_*ij*_, where *j* iterates through all nodes that share an element with node ***n***_*i*_. We then solve our nodal system for mechanical equilibrium such that the total force acting on every node is zero (i.e. ***F***(***n***)=0). In keeping with previous methodologies we assume that inertial forces have no significant impact on the system (46). Due to the non-linearity of the equations, we use a Newton-Raphson numerical scheme to attain a solution (66).

We first consider a system of *m* cytoskeletal elements *ε*_*cγto*_ connected to a fixed node at the cell center (Figure S5); in this study *m*=21. The elements have a reference length equal to the initial radius of the cell *R*_0_ and are> aligned in the radial direction, equally separated at an angle *ω*=30*°/*(*m* - 1). The passive cytoskeletal stiffness *k*_*ec*_ may be *c*onverted to a spring stiffness via *k*_*ec*_=*k*_*ec*_*A*_0_*/R*_0_, where *A*_0_ is a reference cross-sectional area assumed to be constant for all elements. This force acts in parallel with the active cytoskeletal force. Contractility *ρ*_*ij*_ and polymerization-induced stress *σ*_p,*ij*_ are modeled as described by Eqns 3 and 5, with the element strain given by 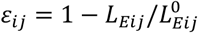. At any given time point, we can compute the active force such that 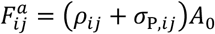, with the total cytoskeletal element force given by 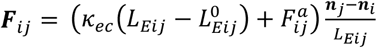, if *E_ij_* ∈ ***E***_*cryto*_. These elements are constrained to displace only in the radial direction (*u*_*θ*_=0). The outer cytoskeletal nodes are connected by membrane elements ***E***_*mem*_, whose passive forces are similarly described by 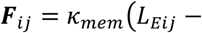, if *E_ij_* ∈ ***E***_*mem*_. Adhesions form and fail between nodes on membrane and those on the neighboring cell. To describe this activity, we again take advantage of symmetry and consider there to be a number of fixed nodes along the hexagonal boundary with which a cell-cell adhesion element ***E***_*adh*_ may form (described in more detail in the next section). The effective stiffness of these junction elements is given by *k*_*J,ij*_=*k*_*b*_*c*_*b,ij*_*A*_0_, such that the associated element force is 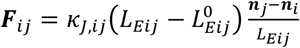, if *E_ij_* ∈ ***E***_*adh*_. It should be noted that when the membrane nodes are disconnected the element stiffness tends to zero (i.e. *k*_*J,ij*_ *→ 0)*.

### Simulation procedure

The model is implemented using an ODE solver (ode15s) in Matlab (R2020b, Mathworks). An event function is defined in the option structure of ode15s to identify (i) when a node on the cell membrane is within contact distance of a corresponding node on a neighboring cell membrane, or (ii) when the force in adherens junction bonds exceeds a critical rupture force. This event respectively triggers the formation of adhesive bonds between the cells described by an element with stiffness *k*_*J,ij*_=*k*_*b*_*c*_*b,ij*_*A*_0_ and reference length *L*_*J*_, or adhesion failure via the element stiffness tending to zero (*k*_*J,ij*_ → 0). The simulation proceeds as follows:

1. Initialize a symmetric cell of radius *R*_0_ with minimum gap distance *L*_*g*_ from its neighbor (Figure S5).
2. Solve the network force balance and evolution of cytoskeletal dynamics over time.
3. At every time point assess if the position of any membrane node ***n***_*i*_ *∈ **n***_*mem*_ is within contact distance of a node on the neighboring cell (*L*_*g,i*_ *≤ L*_]_). If true, activate an adhesion element (*k*_*J,ij*_=*k*_*b*_*c*_*b,ij*_*A*_0_).
4. At every time point assess if the cadherin bond force within any adhesion element ***E***_*ij*_ *∈ **E***_*adh*_ has exceeded the critical rupture force (*F*_*b,ij*_ *≥ F*_*crit*_). If true, remove the adhesion element (*k*_*J,ij*_ *→ 0)*.
5. When the simulation time reaches *t*_*max*_ end the simulation.

By combining and solving the ordinary differential equations (Eqns 3,5) in conjunction with network force balance, we attain a numerical solution for endothelial monolayer dynamics.

## Acknowledgments

This work was supported by NIH Awards U01 CA202177 (R.D.K, V.S.) and R01GM095977 (T.M.S); NCI Awards R01CA232256 (V.B.S.), R01CA232256 (A.T.W.) and R01CA207935 (A.T.W.); NSF CEMB Grant CMMI-154857 (V.B.S.); NSF Grants MRSEC/DMR-1720530 and DMS-1953572 (V.B.S.); Cancer Research UK Multidisciplinary Award C57744/A22057 (E.M.); UKRI Future Leaders Fellowship MR/T043571/1 (F.S.); Spanish Ministry of Science, Innovation and Universities RTI2018-094494-B-C21 (J.M.G.A.); NIBIB Awards R01EB017753 and R01EB030876 (V.B.S.)

## Contributions

E.Mc. and V.B.S. designed the theoretical models; E.Mc. and T.S. carried out the computations; X.C. developed an analytical model; E.M., G.E.M., A.T.M., T.M.S, and R.D.K. designed and conducted the experiments; E.Mc., E.M., T.M.S., R.K. and V.B.S. analysed and interpreted the data; E.Mc., T.S., and V.B.S. wrote the manuscript. All authors contributed to manuscript editing.

## Supplementary Information

### SI Section S1: Active transport of cadherin (exo-/endocytosis)

The density of unbound cadherin *c*_*u*_ at a junction site is not necessarily fixed, but can change as dependent on active transport processes. Cadherin proteins are transported from within the cell to the membrane through exocytosis, a process we may consider to occur at rate *k*_*exo*_. Conversely, endocytosis is the process of moving materials from the membrane into the cell (67), occurring at rate *k*_*endo*_. Initially neglecting cell-cell contact phenomena, a simple kinetic equation to describe the rate of cadherin transport to/from a junction may be given by:

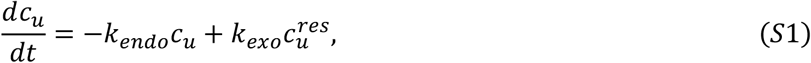

where 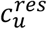 is the cadherin concentration in some reservoir. While exocytosis is considered to operate continuously (68), active transport from the membrane is known to be mechanosensitive; low junction tension promotes cadherin loss by endocytosis (49). Thus, we may consider the endocytosis rate coefficient to be stress-dependent such that:

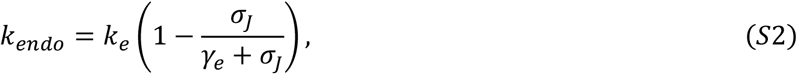

where parameter *γ*_*e*_ *controls* the sensitivity of endocytosis processes to junction stress *σ* _*J*_. For illustration purposes we can simplify further by assuming a single governing rate for the transport processes *k*_*exo*_=*k*_*e*_, leading to a reduced form:

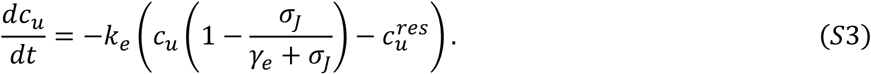

With addition of the catch/slip model and main framework, the change in the density of unbound cadherin at a junction can be expressed as:

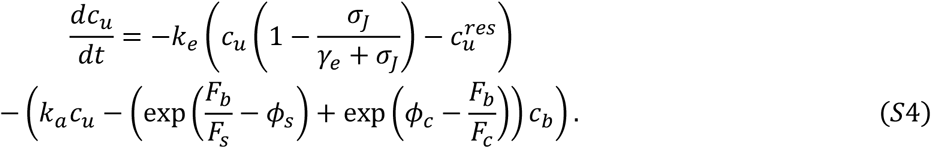

**Figure S1:**
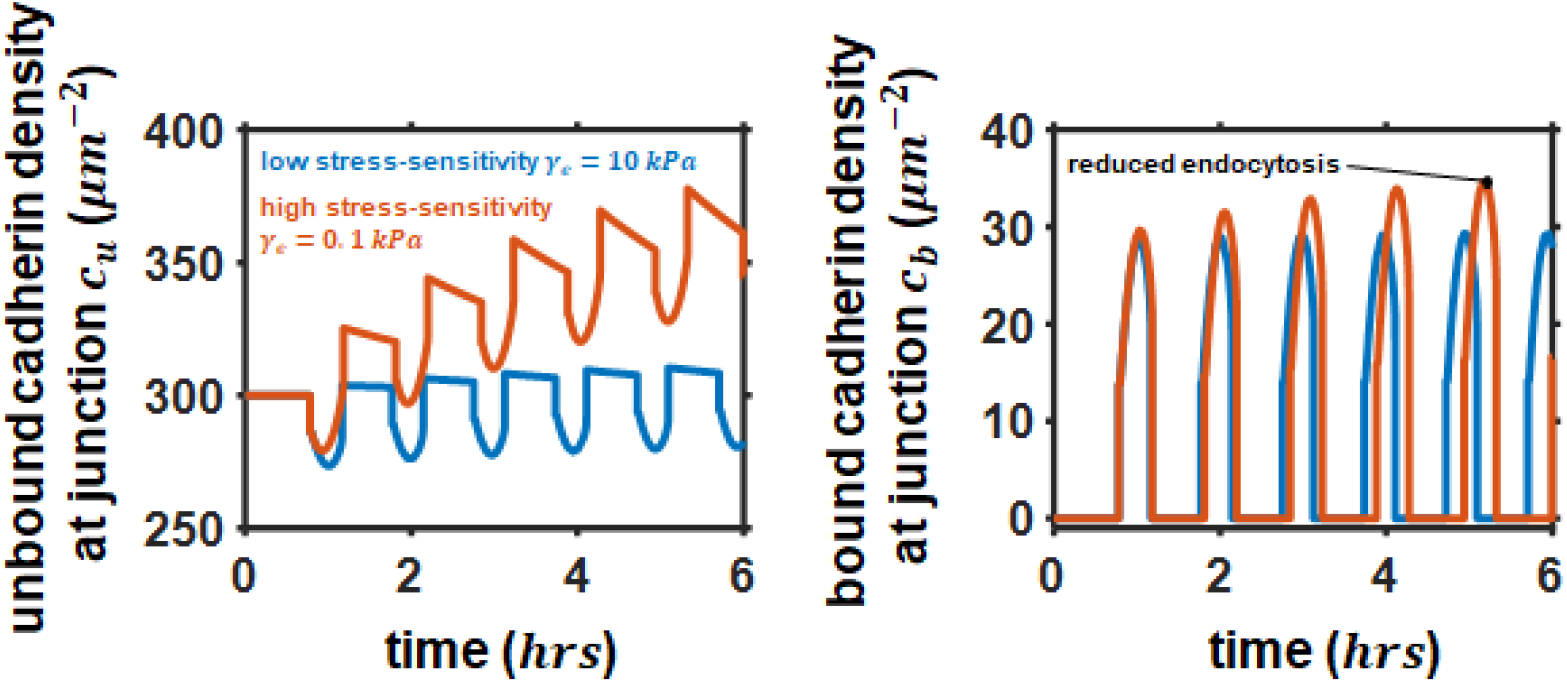
Predicted unbound and bound density of cadherin within a cell-cell adhesion over a number of cycles with either low sensitivity of endocytosis to stress (*γ*_*e*_=10 *kPa*), or a high sensitivity *γ*_*e*_ *=*0.1*kPa* For these simulations *α*_*c*_=20*kPa*^*−1*^, *P*_0_=− 3. 75 *kPa,k*_*e*_*=*1×*10*^*−4*^ *s*^*−1*^, and 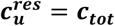.

Here, by changing the sensitivity parameter *γ*_*e*_ we can regulate the balance between exo- and endocytosis for a junction site over time as dependent on junction stress *σ*_*J*_ (Figure S1). Without low stress sensitivity, we see that the maximum cadherin density remains effectively constant during each “rupture and heal” cycle. Reducing *γ*_*e*_ increases the sensitivity of endocytosis to stress, thereby promoting increased cadherin protein reinforcement under tension by reducing endocytosis (while exocytosis operates continuously at a fixed rate).

It is also worth noting that here we implicitly assume a constant reservoir concentration as we consider the number of unbound cadherin proteins in the reservoir is significantly higher that at the junction (i.e. 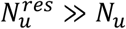) such that transport has a negligible influence on the cytosolic/reservoir concentration. However, when applying the framework to a large number of junctions the total number of cadherin proteins 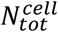 should be carefully conserved in line with our previous work on focal adhesions (69) by enforcing 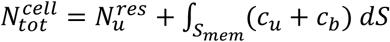. Here 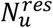 is the number of cadherin proteins within the cell body and the second term determines the total number in the membrane (of area *S*_*mem*_).

### SI Section S2: Derivation of 1D equilibrium equations

The cell stress depends on both active and passive constituents, such that *σ* _*ec*_=*ρ + k*_*ec*_ *ε* _*ec*_ where *K*_*ec*_ is the effective passive cytoskeletal stiffness. Initially, considering a cellular length of interest *L*_*ec*_ and assuming the reference state to be a stress-free connected junction, we can derive an expression for the system stress as governed by mechanical equilibrium. As shown in Figure 1B, we envisage the cell and junction as elements in series; this indicates that the stress in the cell and junction is equivalent, i.e. *σ* _*tot*_= *σ* _*ec*_= *σ* _*J*_, where *σ* _*J*_ is the stress acting on the junction. Similarly, assuming symmetry in the connected cells, the total change in the length of our system is zero such that *L*_*ec*_*ε*_*ec*_ *+ L*_]_ *ε* _*j*_*=0*, where *L*_*J*_ is the reference junction length and *ε*_*J*_ the junction strain. This strain will be governed by the effective junction stiffness (*K*_*j*_=*k*_*b*_*c*_*b*_*L*_*J*_), whereby *ε*_*J*_= *σ*_*tot*_*/*(*k*_*b*_*c*_*b*_*L*_*J*_). We can thus derive an equilibrium expression via:

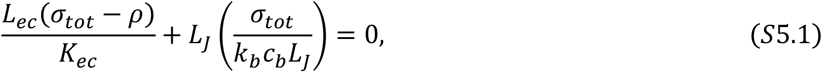

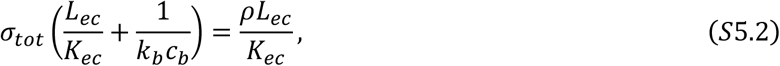

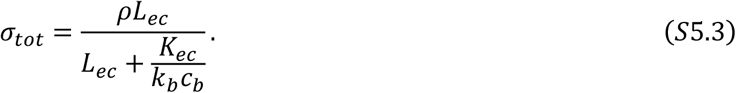

As we progress to consider the role of actin polymerization and associated polymerization-induced stress *σ* _P_ in junction recovery, we evolve this expression (Eqn S5.3) to a reference state with disconnected cells separated by a gap of length *L*_*g*_. The change in cell and junction length therefore leads to a boundary condition *L*_*ec*_*ε*_*ec*_ *+ L*_]_ *ε* _]_*=L*_*g*_. Assuming the polymerization-induced stress acts in parallel with the cell passive stress and contractile stress, we expand Eqn S5.1 to find:

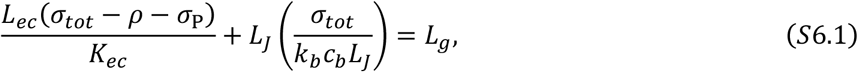

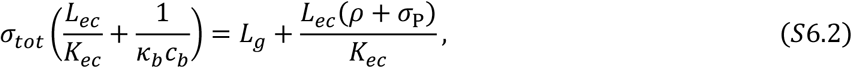

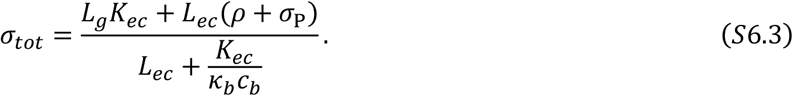

We note that due to sign convention fro compression, the value of *σ*_*P*_ will be negative 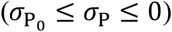.

### SI Section S3: An anlytical analysis of the stress concentration at triple-cell junctions

We consider a triple-cell junction with a geometry as shown by Figure S2A. We assume both the cell cytoskeleton and adhesion adopt angular shapes such that the polar coordinate (*r, θ*) can be conveniently applied. The cell sectors are interconnected with adhesion sectors (represented by boundary lines)(70). For illustration purposes, we assume that both the cell and adhesions are linear elastic. In either sector, the stress fields *σ* _*r*_, *σ* _*θ*_, *τ* _*r θ*_ and the displacement fields *u*_*r*_, *u*_*θ*_ in the cells or the adhesions should follow (71):

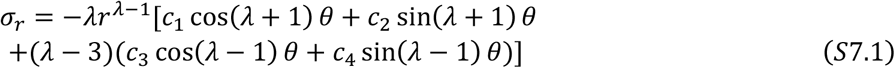

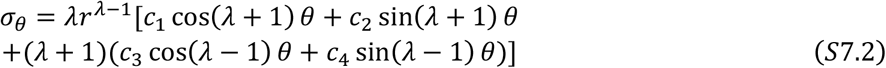

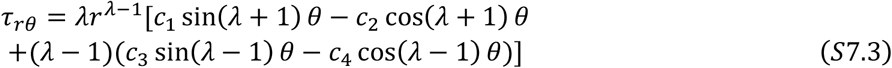

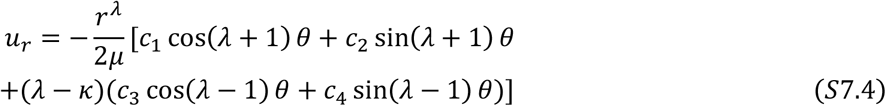

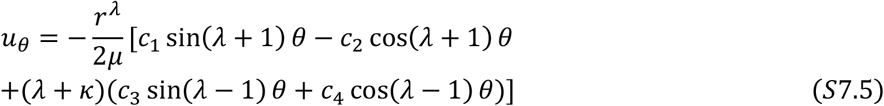

Here, *μ* is the shear modulus and *k*=(3 *- *v*)/(1* + *v*), *v* being the Poisson’s ratio. *c*_*i*_(*i*=1,2,3,4) are constants and *λ* is a variable (70) that controls how stresses scale with the distance to the junction. Note that the mechanical properties (*μ, v*) and different sets of coefficients *c*_*i*_(*i*=1,2,3,4) will be different for either the cell or adhesion sector. At the cell-adhesion interface, the stress and displacement must satisfy the continuity condition

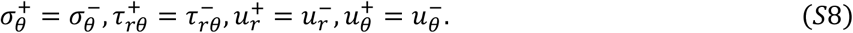

Here the superscripts + and – *de* note the two sides of the cell-adhesion boundary. With the representation of displacement and stresses, the singularity at the cell junction can be investigated. We consider a case where three cells form a triple cell junction with 3-fold symmetry (*α*=2*°, ϕ* =118*°*) as shown by Figure S2A. By taking advantage of the symmetry, the problem is reduced to Figure S2B. To satisfy the symmetry, on boundary 1 and 2, *u*_0_=0, *τ* _*r θ*_ *=0*. Combining the boundary conditions with Eqns S1-S5, we obtain a homogeneous linear equation

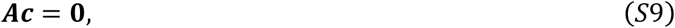

where the vector 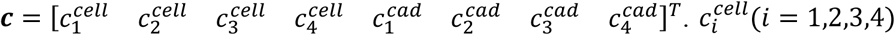. and 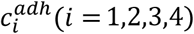 denote the four undetermined constants *c*_*i*_ in Eqns S1-S5 for the cell and adhesion, respectively. Defining 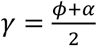, the 8 by 8 matrix ***A*** can be written as

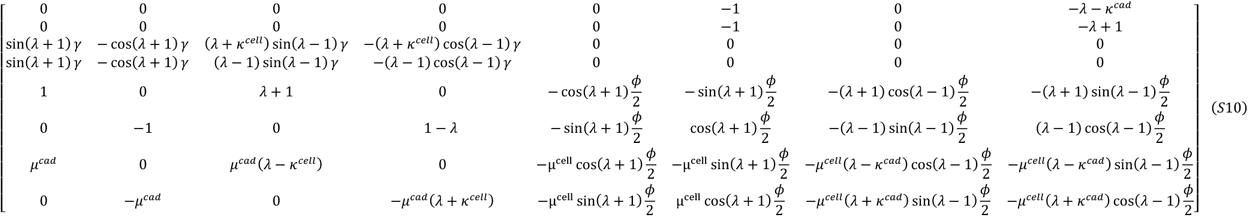

Since Eqn S9 is homogeneous, for there to be a non-trivial solution ***c***, the determinant of ***A*** must vanish (|***A***|=0). Solving this equation for the typical parameters given in Table S1, we find *λ -* 1*=-*0.37 given the typical properties listed in Table S1. Note that *σ*_*r*_⍰*r*^*λ− 1*^ and *σ* _*θ*_⍰*r*^*λ− 1*^ both scale with *r*^*λ − 1*^, therefore a value of 0 *< λ < 1* results in infinitely large stresses near the junction as *r →* 0 (Eqns S7.1-S7.3). This result shows that a symmetric triple cell junction is inherently unstable with a stress singularity at the junction.

**Table S1.** Cell and adhesion mechanical properties.

| Symbol | Physical meaning | Value |
| --- | --- | --- |
| $\mu^{cell}$ | Cell shear modulus | 1kPa |
| $\mu^{adh}$ | Adhesion effective shear modulus | 0.01kPa |
| $\nu^{cell}$ | Cell Poisson's ratio | 0.3 |
| $\nu^{adh}$ | Adhesion effective Poisson's ratio | 0.3 |

**Figure S2:**
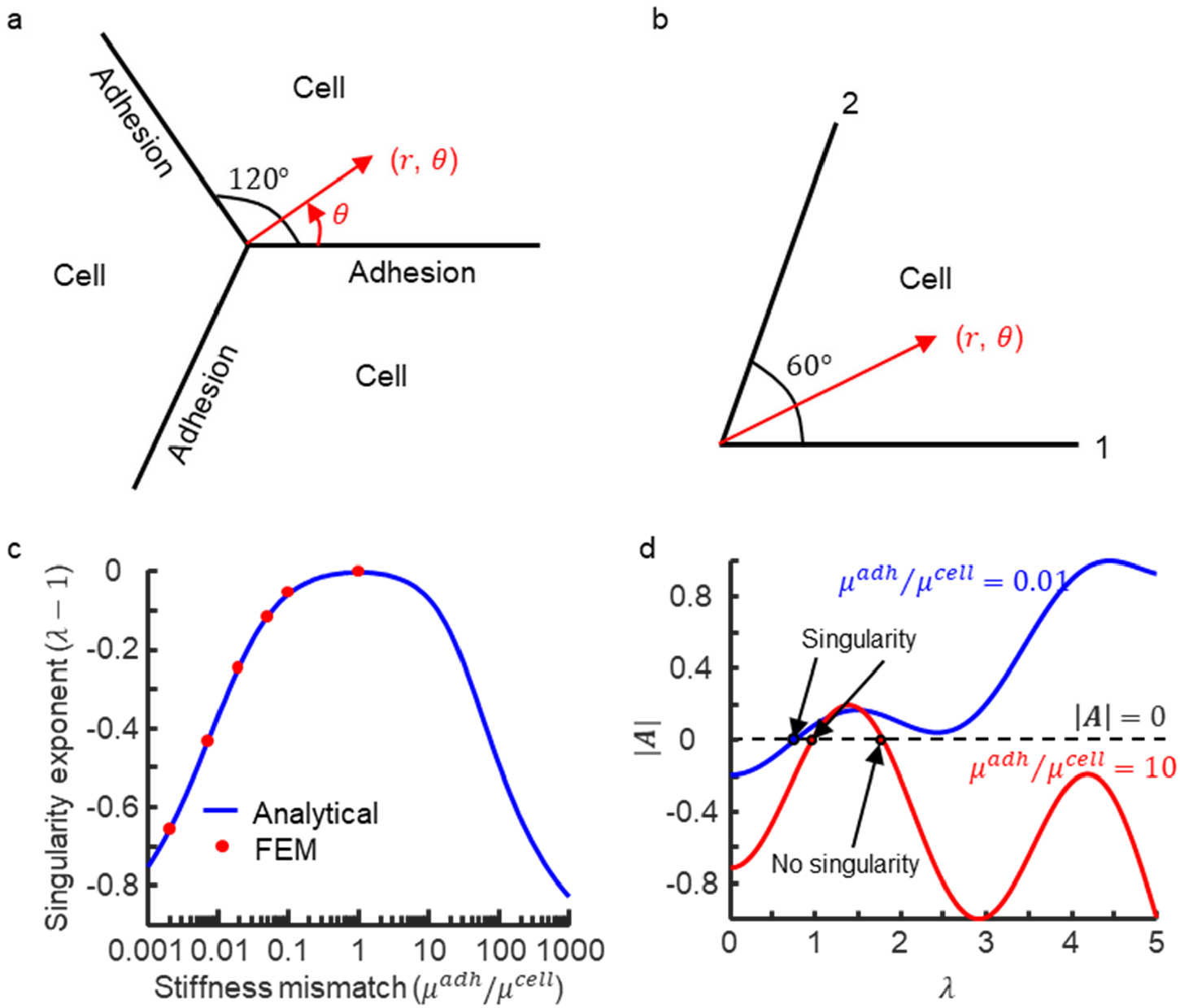
A) The triple cell junction geometry. We assume that around the junction the cell and cadherin-mediated adhesion follow a 3-fold symmetry. The cell spans an angle of *ϕ =*118° and the adhesions span an angle of *α=*2° B) The repeating unit of the triple cell junction; C) The minimum singularity exponent as a function of the adhesion-cell stiffness mismatch. The singularity vanishes when the adhesion and the cells have the same stiffness (*μ*^*adh*^*/ μ*^*cell*^*=*1); D) |*A*| as a function of *λ* for *μ*^*adh*^ */ μ* ^*cell*^*=*0.01o*r* 10. A *λ >*1 solution can also be found for*μ*^*adh*^ */μ* ^*cell*^*=*10, indicating that the singularity may not occur.

We further explore how *λ* can be influenced by the stiffness of cells and adhesions by finding the root of |*A*|=0 as we vary the ratio *μ*^*adh*^*/μ*^*cell*^. As shown in Figure S2, we find that the minimum root to |*A*|=0 always lies in the range 0 < λ < 1 when the adhesions and the cells have stiffness mismatch (*μ*^*adh*^*/μ*^*cell*^ *< 1). Th*e stresses become more singular as the *μ*^*adh*^*/μ*^*cell*^ ratio deviates from 1 (Figure S2C). Note when *μ*^*adh*^*/μ*^*cell*^ *> 1*, |*A*|=0 has solutions with *λ > 1 (71)*, which will not introduce a singularity (Figure S2D). In such case, only with appropriate loading and boundary conditions, can the singularity arise. To further confirm whether the singularity can occur when there is stiffness mismatch (*μ*^*adh*^*/μ*^*cell*^ *< 1)*, we built finite element models (FEM) and assessed the stress near the triple-cell junction. When we apply an isotropic contractile stress that mimics an active endothelial cell contraction, we find that the singularity arises at the triple-cell junction. The singularity exponent obtained from FEM agrees excellently with the analytical analysis for *μ*^*adh*^*/μ*^*cell*^ *< 1 (Fig*ure S2C). These results suggest that the stress singularity was due to the mismatch of the cell and adhesion moduli.

### SI Section S4: Model parameters

**Table S2:**
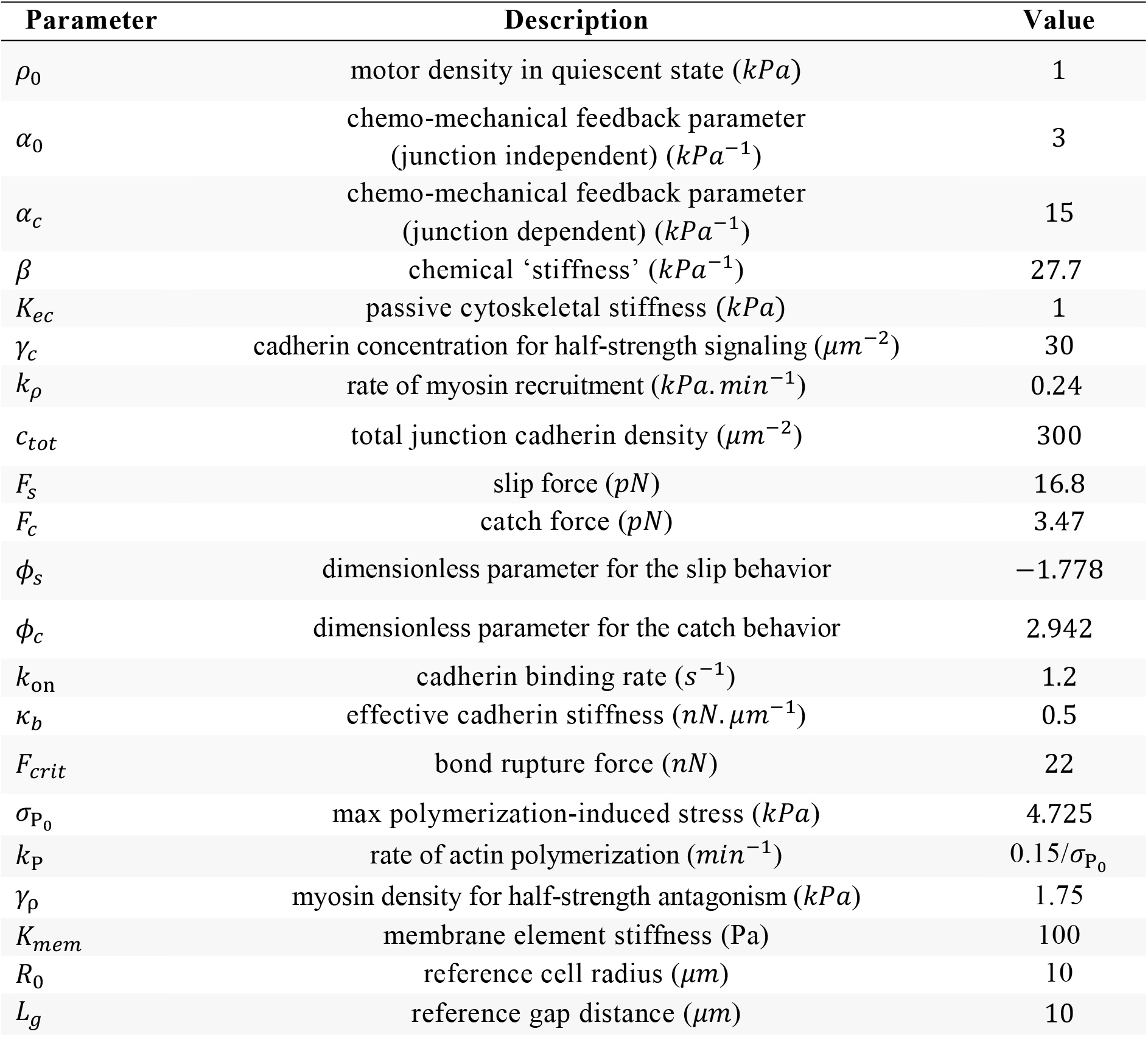
Parameters for chemo-mechanical model.

| Parameter | Description | Value |
| --- | --- | --- |
| $\rho_0$ | motor density in quiescent state ($kPa$) | 1 |
| $\alpha_0$ | chemo-mechanical feedback parameter<br>(junction independent) ($kPa^{-1}$) | 3 |
| $\alpha_c$ | chemo-mechanical feedback parameter<br>(junction dependent) ($kPa^{-1}$) | 15 |
| $\beta$ | chemical ‘stiffness’ ($kPa^{-1}$) | 27.7 |
| $K_{ec}$ | passive cytoskeletal stiffness ($kPa$) | 1 |
| $\gamma_c$ | cadherin concentration for half-strength signaling ($\mu m^{-2}$) | 30 |
| $k_\rho$ | rate of myosin recruitment ($kPa \cdot min^{-1}$) | 0.24 |
| $c_{tot}$ | total junction cadherin density ($\mu m^{-2}$) | 300 |
| $F_s$ | slip force ($pN$) | 16.8 |
| $F_c$ | catch force ($pN$) | 3.47 |
| $\phi_s$ | dimensionless parameter for the slip behavior | -1.778 |
| $\phi_c$ | dimensionless parameter for the catch behavior | 2.942 |
| $k_{on}$ | cadherin binding rate ($s^{-1}$) | 1.2 |
| $\kappa_b$ | effective cadherin stiffness ($nN \cdot \mu m^{-1}$) | 0.5 |
| $F_{crit}$ | bond rupture force ($nN$) | 22 |
| $\sigma_{P_0}$ | max polymerization-induced stress ($kPa$) | 4.725 |
| $k_P$ | rate of actin polymerization ($min^{-1}$) | $0.15/\sigma_{P_0}$ |
| $\gamma_\rho$ | myosin density for half-strength antagonism ($kPa$) | 1.75 |
| $K_{mem}$ | membrane element stiffness (Pa) | 100 |
| $R_0$ | reference cell radius ($\mu m$) | 10 |
| $L_g$ | reference gap distance ($\mu m$) | 10 |

The parameters associated with cytoskeletal contractility are confined to previously reported ranges (33, 72); namely the quiescent myosin density *ρ*_0_=1 *kPa, f3=*27.7 *kPa* where 0 *< (α*_*c*_ *+ α*_0_)*/β < 1, and K*_*ec*_=1 *kPa*. The density of cadherin proteins at adherens junctions has been measured as on the order of 700 *µm*^*2*^ *(73*), motivating us to confine *c*_*tot*_ to a ma*g*nitude of 300 *- 500 µm*^*2*^. Catch-slip model parameters are estimated from the experimental data of Manibog *et al*. (30), as shown in Figure 1C such that *F*_*s*_=16.8 *pN, F*_*c*_=3.47 *pN, ϕ*_*s*_*=-1*.778, *ϕ*_*c*_=2.942, and *k*_*b*_=0.5 *nN. µm* ^*1*^. Manibog *et al*. (30) also suggest a threshold force of *F*_*crit*_=22 *pN* for the unbinding of a single VE-cadherin bond. The protein binding rate is assumed to be constant and follows Bangasser *et al*. (74) whereby *k*_*on*_=1.2 *s* ^*1*^. We assume a membrane element stiffness of *K*_*mem*_=100 *Pa*. The myosin recruitment and polymerization rates (*k*_*ρ*_, *k*_*P*_) and the feedback parameters (*γ*_*c*_, *γ*_*ρ*_) are calibrated to provide good agreement with experimental timescales (Figure 5), with the influence of polymerization parameters shown in Figure S3A-B.

### SI Sections S5: Additional figures

**Figure S3:**
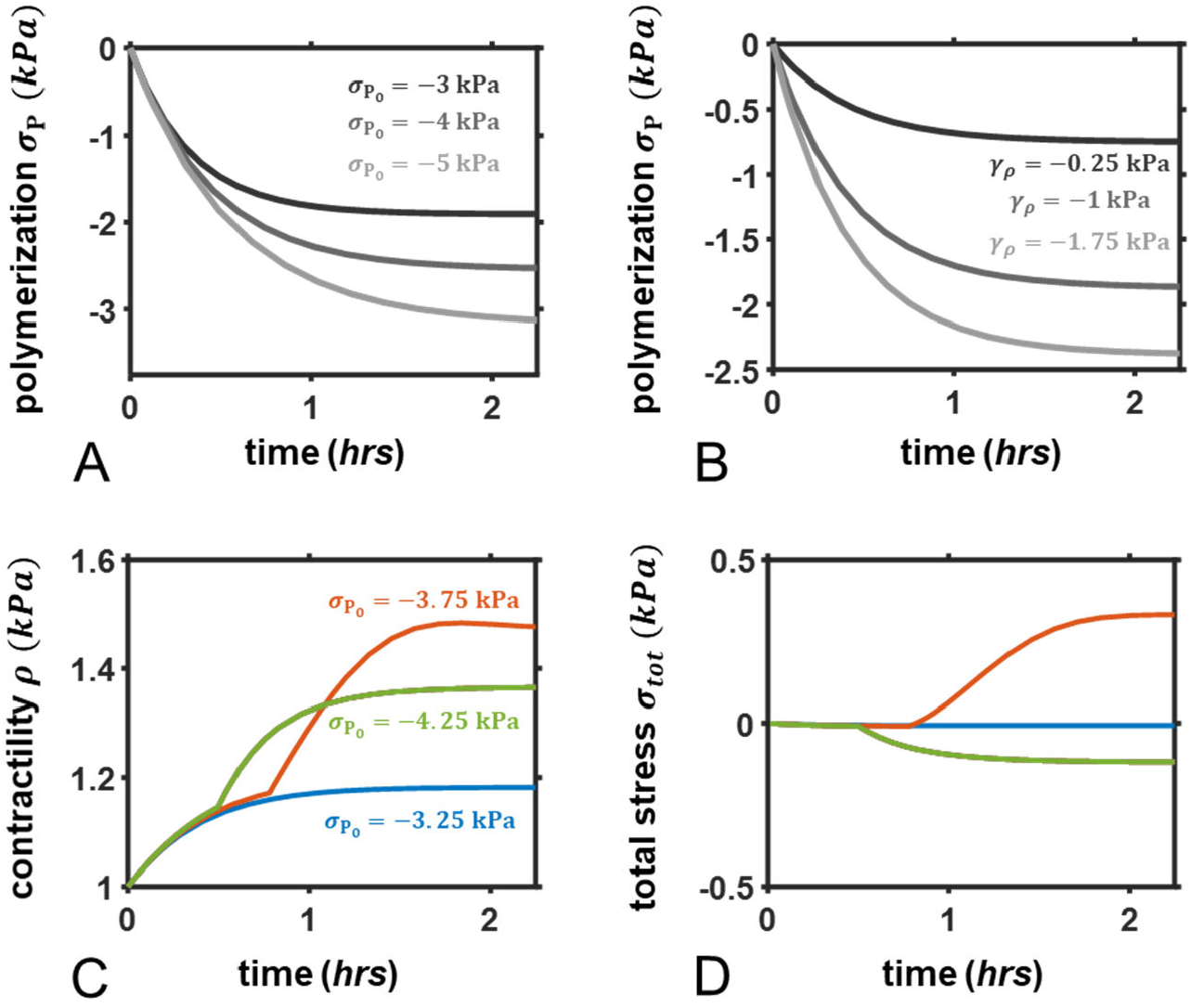
Predicted polymerization-induced stress in response to a change in A) 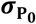 *and B) γ* _*ρ*_ for a fixed contractility *ρ =*1*kPa*; Predicted C) contractility *ρ* and D) total stress *σ*_*tot*_ in response to low, intermediate, and high polymerization.

**Figure S4:**
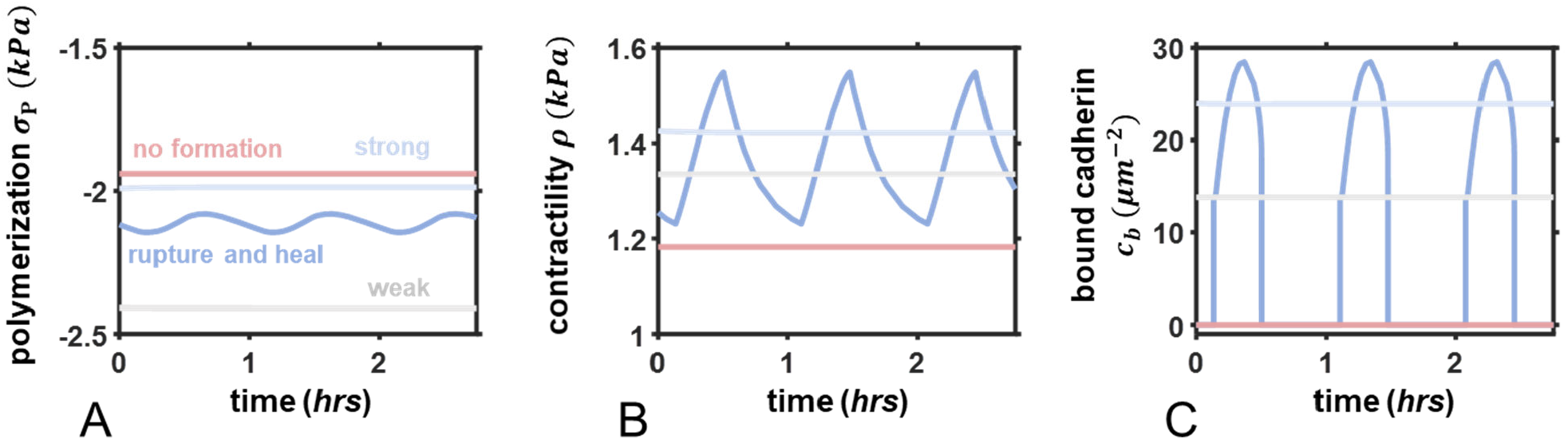
Steady state A) polymerization-induced stress, B) contractility, and C) bound cadherin density in each phase: rupture/heal (*α*_*c*_*=*20*kPa*^*−1*^; 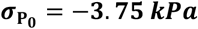), strong (*α*_*c*_=10*kPa*^*−1*^; 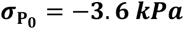), weak (*α*_*c*_=10*kPa*^*−1*^; 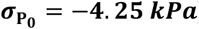), none (*γ*_*c*_=10 *kPa*^*−1*^; 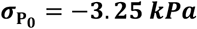).

**Figure S5:**
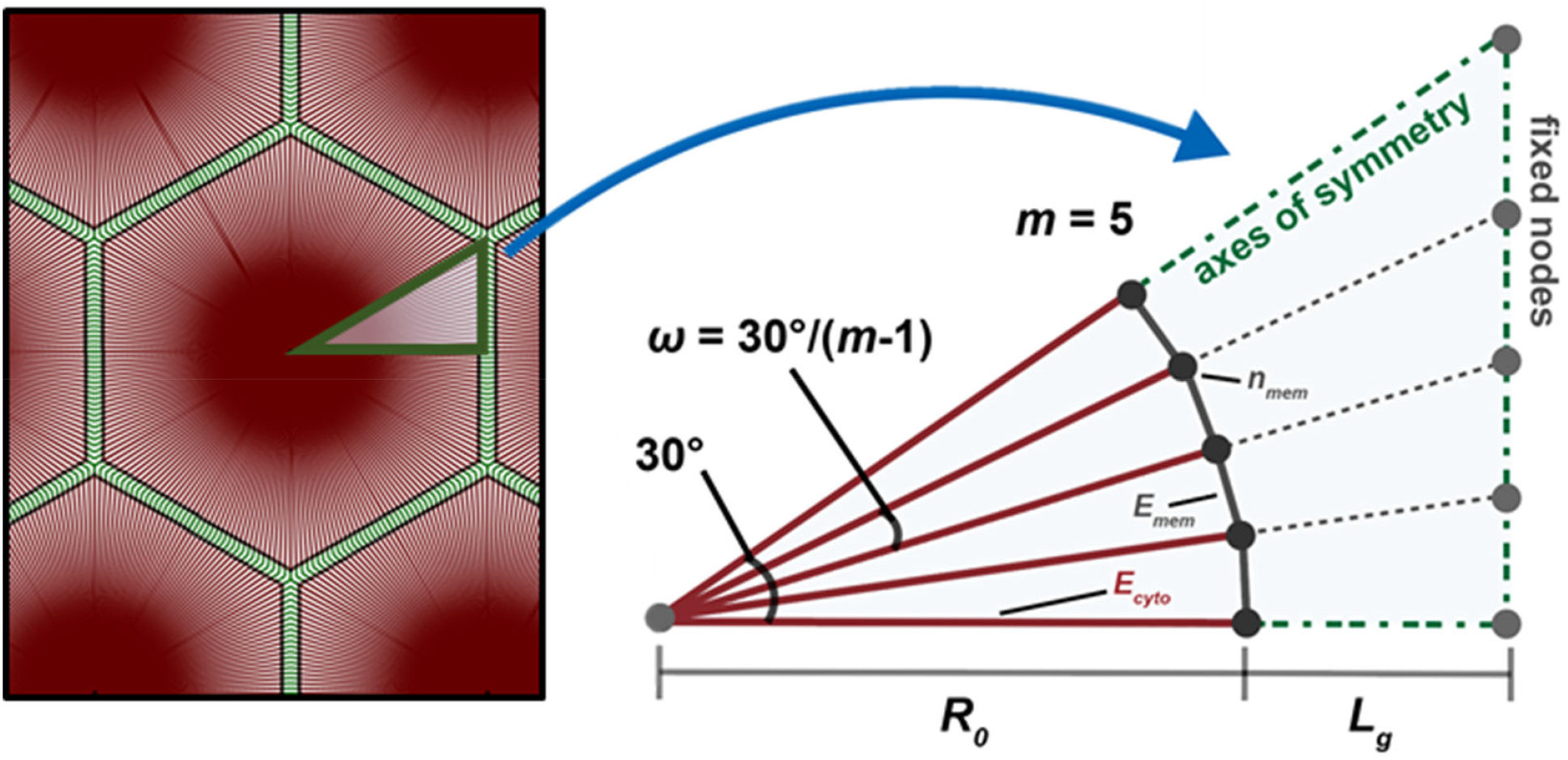
Schematic of symmetric cell geometry in reference state.

## Notes

### Competing Interest Statement

The authors have declared no competing interest.

